# Tethering by Uso1 is dispensable: The Uso1 monomeric globular head domain interacts with SNAREs to maintain viability

**DOI:** 10.1101/2022.11.30.518472

**Authors:** Ignacio Bravo-Plaza, Víctor G. Tagua, Herbert N. Arst, Ana Alonso, Mario Pinar, Begoña Monterroso, Antonio Galindo, Miguel Á. Peñalva

**Affiliations:** Departments of Cellular and Molecular Biology, CSIC Centro de Investigaciones Biológicas, Ramiro de Maeztu 9, 28040 Madrid Spain; Instituto de Tecnologías Biomédicas, Hospital Universitario Nuestra Señora de Candelaria, Santa Cruz de Tenerife, Spain.; Department of Infectious Diseases, Faculty of Medicine, Flowers Building, Imperial College, Armstrong Road, London, SW7 2AZ, UK; Departments of Structural and Chemical Biology, CSIC Centro de Investigaciones Biológicas, Ramiro de Maeztu 9, 28040 Madrid Spain; Division of Cell Biology, MRC Laboratory of Molecular Biology, Francis Crick Avenue, Cambridge, CB2 0QH, UK.

## Abstract

Uso1/p115 and RAB1 tether ER-derived vesicles to the Golgi. Uso1/p115 contains a globular-head-domain (GHD), a coiled-coil (CC) mediating dimerization/tethering and a C-terminal region (CTR) interacting with golgins. Uso1/p115 is recruited to vesicles by RAB1. Paradoxically, genetic studies placed Uso1 acting upstream of, or in conjunction with RAB1 (Sapperstein et al., 1996). We selected two missense mutations in *uso1* resulting in E6K and G540S substitutions in the GHD permitting growth of otherwise inviable *rab1-*deficient *Aspergillus nidulans.* Remarkably, the double mutant suppresses the complete absence of RAB1. Full-length Uso1 and CTRΔ proteins are dimeric and the GHD lacking the CC/CTR is monomeric irrespective of whether they carry or not E6K/G540S. Microscopy showed recurrence of Uso1 on puncta (60 sec half-life) colocalizing with RAB1 and less so with early Golgi markers Sed5 and GeaA/Gea1/Gea2. Localization of Uso1 but not of Uso1^E6K/G540S^ to puncta is abolished by compromising RAB1 function, indicating that E6K/G540S creates interactions bypassing RAB1. By S-tag-coprecipitation we demonstrate that Uso1 is an associate of the Sed5/Bos1/Bet1/Sec22 SNARE complex zippering vesicles with the Golgi, with Uso1^E6K/G540S^ showing stronger association. Bos1 and Bet1 bind the Uso1 GHD directly, but Bet1 is a strong E6K/G540S-independent binder, whereas Bos1 is weaker but becomes as strong as Bet1 when the GHD carries E6K/G540S. AlphaFold2 predicts that G540S actually increases binding of GHD to the Bos1 Habc domain. In contrast, E6K seemingly increases membrane targeting of an N-terminal amphipathic *α*-helix, explaining phenotypic additivity. Overexpression of E6K/G540S and wild-type GHD complemented *uso1Δ*. Thus, a GHD monomer provides the essential Uso1 functions, demonstrating that long-range tethering activity is dispensable. Therefore, when enhanced by E6K/G540S, Uso1 binding to Bos1/Bet1 required to regulate SNAREs bypasses both the contribution of RAB1 to Uso1 recruitment and the reported role of RAB1 in SNARE complex formation (Lupashin and Waters, 1997), suggesting that the latter is consequence of the former.

## Introduction

Vesicular traffic at the ER/Golgi interface is the cornerstone of the secretory pathway (Barlowe and Miller, 2013; Weigel et al., 2021). In current models, in which traffic across the Golgi is driven by cisternal maturation (Day et al., 2013; Pantazopoulou and Glick, 2019), COPII vesicles generated at specialized domains of the ER fuse homotypically and heterotypically to form and feed the earliest Golgi cisternae (Rexach et al., 1994). As straightforward as this step might seem, it involves a sophisticated circuitry of regulation. Actual fusion is in part mediated by compartmental-specific sets of four-membered SNARE protein complexes (SNARE bundles) (Malsam and Sollner, 2011; Pelham, 2001; Rizo and Sudhof, 2012). Most SNARES are type II single TMD proteins, whose N-terminal cytosolic domain contains nearly all the polypeptide, excepting a few lumenal residues. Like any other transmembrane proteins, SNAREs are synthesized in the ER. This implies that they have to travel to compartments of the cell as distant as the plasma membrane in a conformation that precludes them of catalyzing what would be a calamitous fusion of non-cognate donor and acceptor compartments. Achieving the strictest specificity is particularly challenging in the ER-to-Golgi stage that, as the first step in the secretory pathway, represents an obligate point of transit for each and every transmembrane SNARE. Therefore, the only SNARES acting in this first step are the Qa Sed5, the Qb Bos1, the Qc Bet1 and the R-SNARE Sec22, which form the bundle mediating fusion of carriers that coalesce into cisternae (McNew et al., 2000).

Given the central role played by the secretory pathway in the physiology of every eukaryotic cell, it is unsurprising that this step involves regulatory factors which are essential for cell survival. One is the SM (<u>S</u>ec1, Munc-18) protein Sly1, which promotes SNARE bundle formation (Bracher and Weissenhorn, 2002; Peng and Gallwitz, 2002; Thomas et al., 2019) Another is the TRAPPIII complex, which interacts with the external coat of COPII carriers and acts as a guanine nucleotide exchange factor (GEF) for RAB1 (Bracher and Weissenhorn, 2002; Cai et al., 2007; Galindo et al., 2021; Joiner et al., 2021; Lord et al., 2011; Peng and Gallwitz, 2002; Pinar and Peñalva, 2020; Riedel et al., 2017; Thomas et al., 2018; Thomas et al., 2019). This small GTPase is a key player that transiently recruits protein effectors from the cytosol to donor and acceptor membranes (Sogaard et al., 1994) and regulates SNARE assembly through an as yet undefined mechanism (Lupashin and Waters, 1997; Sapperstein et al., 1996). One RAB1 effector is a fungal protein denoted Uso1, whose highly conserved metazoan homologue is p115. These are homodimers with a globular N-terminal head and a long C-terminal coiled-coil region characteristic of tethering proteins, which bring donor and acceptor membranes into the distance at which v- and t-SNAREs can engage into the productive *trans*-SNARE complex that mediates membrane fusion (Cao et al., 1998; Nakajima et al., 1991; Sapperstein et al., 1996; Sapperstein et al., 1995; Seog et al., 1994; Yamakawa et al., 1996).

Despite most Golgi tethers are functionally redundant, Uso1 is unique in that it is an essential protein. *uso1-1*, a *S. cerevisiae* amber mutation truncating most, but not all the coiled-coil region, is viable, yet further upstream truncation removing the complete coiled-coil is lethal, which was taken as evidence that tethering is the essential function of Uso1 (Seog et al., 1994). In addition, as the coiled-coil is predicted to mediate dimerization, it is broadly accepted that Uso1 is ‘just’ an essential homodimer that tethers vesicles to the acceptor membrane. However, genetic evidence stubbornly indicates that Uso1 plays additional functions related with SNAREs. For example, Sapperstein et al showed that SNAREs function downstream of Uso1 (Sapperstein et al., 1996). Notably, the view that Uso1 is a mere RAB1 effector was challenged by the observation that overexpressing Ypt1 (yeast RAB1) rescues lethality of *uso1Δ*, whereas the reciprocal is not true, indicating that Uso1 acts upstream of or in conjunction with RAB1 (Sapperstein et al., 1996).

Our laboratory is interested in deciphering the domains of action of RAB GTPases in the genetic and cell biological model organism *Aspergillus nidulans* (Pinar and Peñalva, 2021). We have previously gained mechanistic insight into the activation of RAB11 by TRAPPII by exploiting a forward genetic screen for mutations bypassing, at the restrictive temperature, the essential role of the key TRAPPII subunit Trs120 (Pinar et al., 2019; Pinar et al., 2015; Pinar and Peñalva, 2020). In this type of screen, a strain carrying a *ts* mutation in the gene-of-interest is mutagenized and strains bypassing lethality at the restrictive temperature are identified and characterized molecularly. A well-characterized, conditionally lethal *rab1* mutation is available (Pinar et al., 2013), enabling us to investigate pathways collaborating with RAB1 in anterograde traffic. We isolated two *uso1* missense mutations causing substitutions in the globular head domain (GHD). When combined together, these rescued the lethality resulting from *rab1Δ* and promoted the localization of the protein to early Golgi cisternae by increasing Uso1 binding to the cytosolic region of the Qa SNARE Bos1 and, potentially, by improving the interaction of an N-terminal amphipathic *α*-helix with membranes. Importantly, we show that endogenous expression of a protein consisting solely of the double mutant GHD, or overexpression of double mutant or wild-type GHD, rescue the lethality resulting from *uso1Δ*, even though the GHD is monomeric. Our results show that one essential role of RAB1 is recruiting Uso1 to membranes, and that the essential role of Uso1 is not tethering membranes, but rather regulating the formation of the cognate SNARE bundle, indicating that Uso1 is a component of the SNARE fusion machinery.

## Results

### Missense mutations in *uso1* rescue the lethality resulting from *rab1*Δ

*rab1^A136D^* (hereby *rab1^ts^*) mutants do not grow at 37°C. However, when we plated UV-mutagenized conidiospores of the *rab1^ts^* mutant at this temperature, we obtained colonies showing different degrees of growth, presumably carrying mutations rescuing the lethality resulting from *rab1^ts^*. One was chosen for further characterization. By sexual crosses and parasexual genetics this strain was shown to carry a single suppressor mutation, denoted *su1rab1^ts^*, that co-segregated with chromosome VIII. Meiotic mapping narrowed *su1rab1^ts^* to the vicinity (2 cM) of *hisC*. 40 kb centromere distal from *hisC* lies AN0706 (Figure 1A) encoding *Aspergillus nidulans* Uso1, a conserved effector of RAB1. Sanger sequencing revealed the presence of a G16A transition (denoted E6K) resulting in Glu6Lys substitution in the *uso1* gene of *su1rab1^ts^*.

**Figure 1.**
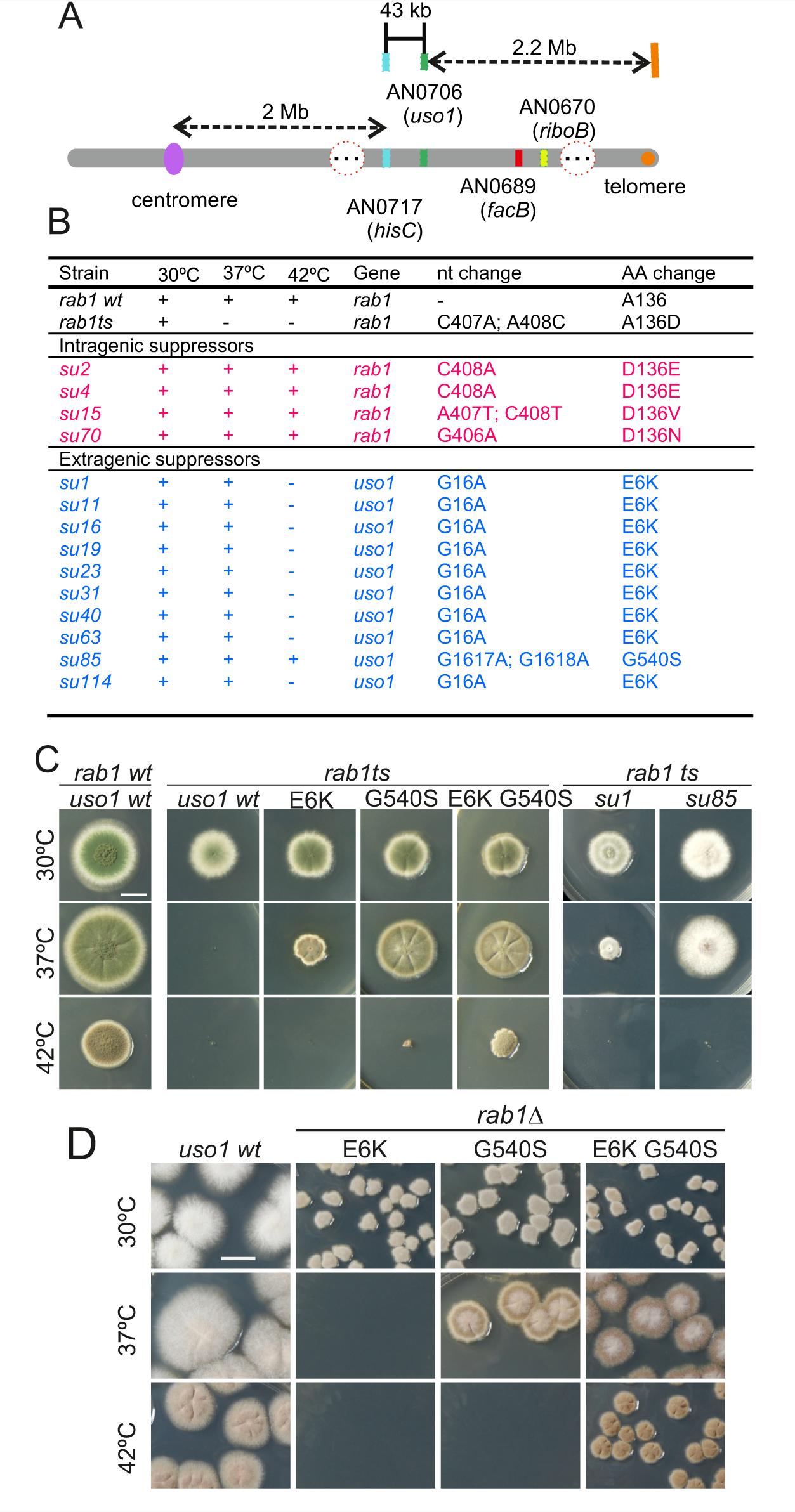
Characterization of mutations bypassing the essential role of RAB1. (A). Genetic map in the region surrounding *uso1* with genetic markers used as landmarks for mapping. (B). Molecular identification of the nucleotide changes in *suArab1ts* strains (C) and (D): growth tests showing *rab1ts*- and *rab1Δ*-rescuing phenotypes, respectively, of individual mutations, and synthetic positive interaction between E6K and G540S. Strains produce either green or white conidiospores (conidiospore colors are used as genetic markers). In (C), strains were point-inoculated. In (D) conidiospores were spread on agar plates to give individual colonies.

To determine if the remaining suppressor strains were allelic to *su1rab1^ts^*, we sequenced *uso1* from a further 13 isolates (Figure 1B). Of these, four were *rab1^ts^* pseudo-revertants that had acquired a functionally acceptable mutation in the altered codon, and eight carried *uso1* E6K, suggesting that the screen was close to saturation. However, one mutation was found to be a different missense allele, *su85rab1ts* (denoted G540S) resulting in Gly540Ser substitution. Single mutant strains carrying these *uso1* mutations showed no growth defect, indicating that E6K and G540S were unlikely to result in loss-of-function, and suggesting instead that mutant strains had acquired features that made them largely independent of RAB1. These findings were unexpected, because in *Saccharomyces cerevisiae* overexpression of Uso1 does not rescue the lethality of *ypt1Δ* mutants (Ypt1 is the yeast RAB1 homologue)(Sapperstein et al., 1996).

To demonstrate that *uso1^E6K^* and *uso1^G540S^* were causative of the suppression, we reconstructed them by homologous recombination. These reverse-genetic alleles rescued viability of *rab1^ts^* at 37°C to a similar extent as *su1rab1^ts^* and *su85rab1^ts^* (Figure 1C). *uso1^G540S^* was the strongest suppressor, such that *rab1^ts^ uso1^G540S^* double mutants grew nearly as the wt at 37°C. Nevertheless, the two alleles showed additivity, and a triple mutant carrying *uso1^E6K^*, *uso1^G540S^* and *rab1^ts^* grew at 42°C, unlike either single mutant (Figure 1C). These data, together with the genetic mapping above, established that *uso1^E6K^* and *uso1^G540S^* are responsible for the suppression phenotype, RAB1 recruits Uso1/p115 to uncoated COPII vesicles and early Golgi cisternae (Allan et al., 2000). Therefore, *uso1^E6K^*^/*G540S*^ might, by increasing the affinity of Uso1 for RAB1, compensate for the reduction in the amount of the GTPase resulting from *rab1^ts^*. However, Figure 1D shows that both *uso1^E6K^ and uso1^G540S^* rescue the lethality resulting from the complete ablation of *rab1Δ* at 30°C, with the strongest *uso1^G540S^* suppressor rescuing viability even at 37°C, and the double mutant rescuing *rab1Δ* even at 42°C (Figure 1D). In contrast, *uso1^E6K^*^/*G540S*^ did not rescue the lethality resulting from *arf1Δ*, nor from *sed5Δ* or *sly1Δ*, the syntaxin and the SM protein which are crucial for the formation of the ER/Golgi SNARE bundle (Figure 1—figure supplement 1), indicating that Uso1 plays a role acting downstream of RAB1 and upstream of or in conjunction with the SNARE machinery. This role is essential for survival (Figure 1—figure supplement 1).

### E6K affects a previously undetected N-terminal helix, whereas G540S is located in a loop near the end of the armadillo domain

1103-residue *A. nidulans* Uso1 is similar in size to 961-residue p115 (bovine) and notably shorter than *S. cerevisiae* Uso1p (1790 residues) (Yamakawa et al., 1996). Thus far, atomic structures of Uso1/p115 are limited to the 600-700 residue GHD, which consists of a highly conserved *α*-catenin-like armadillo-fold (An et al., 2009; Heo et al., 2020; Striegl et al., 2009). *In silico* analyses robustly predict that the approximately C-terminal half of Uso1/p115 consists of a coiled-coil that mediates tethering and dimerization, but this region has not been characterized beyond low resolution EM studies (Yamakawa et al., 1996) . Neither crystal structures nor predictions provided information about the N-terminal extension in which Glu6Lys lies.

Thus, we used AlphaFold2, imposing or not the condition that the protein is a dimer (see below). Figure 2A shows the monomer and dimer models with the highest confidence scores (see Figure 2—figure supplement 1). Like their relatives, Uso1 from *A. nidulans* contains an N-terminal GHD including a previously unnoticed short *α*-helix in which Glu6 affected by E6K lies. This N-terminal extension is followed by ∼34 *α*-helices arranged into 12 tandem repetitions of armadillo repeats (ARM1-ARM12; residues 17 through 564), each containing three right-handed *α*-helices except for the first two repeats. Altogether, the armadillo repeats resemble the shape of a jai alai basket. Downstream of the GHD, AlphaFold2 predicts a long extended coiled-coil (CC) between residues 674 through 1082, which would mediate dimerization (see below) (Figure 2A). The CC ends at a conserved C-terminal region (CTR) (Figure 2A and B), which includes an also C-terminal segment rich in acidic residues. In Uso1, twelve out of the last seventeen amino acids are Asp/Glu (Figure 2A and B). The CC and CTR regions will be collectively denoted the CCD domain.

**Figure 2:**
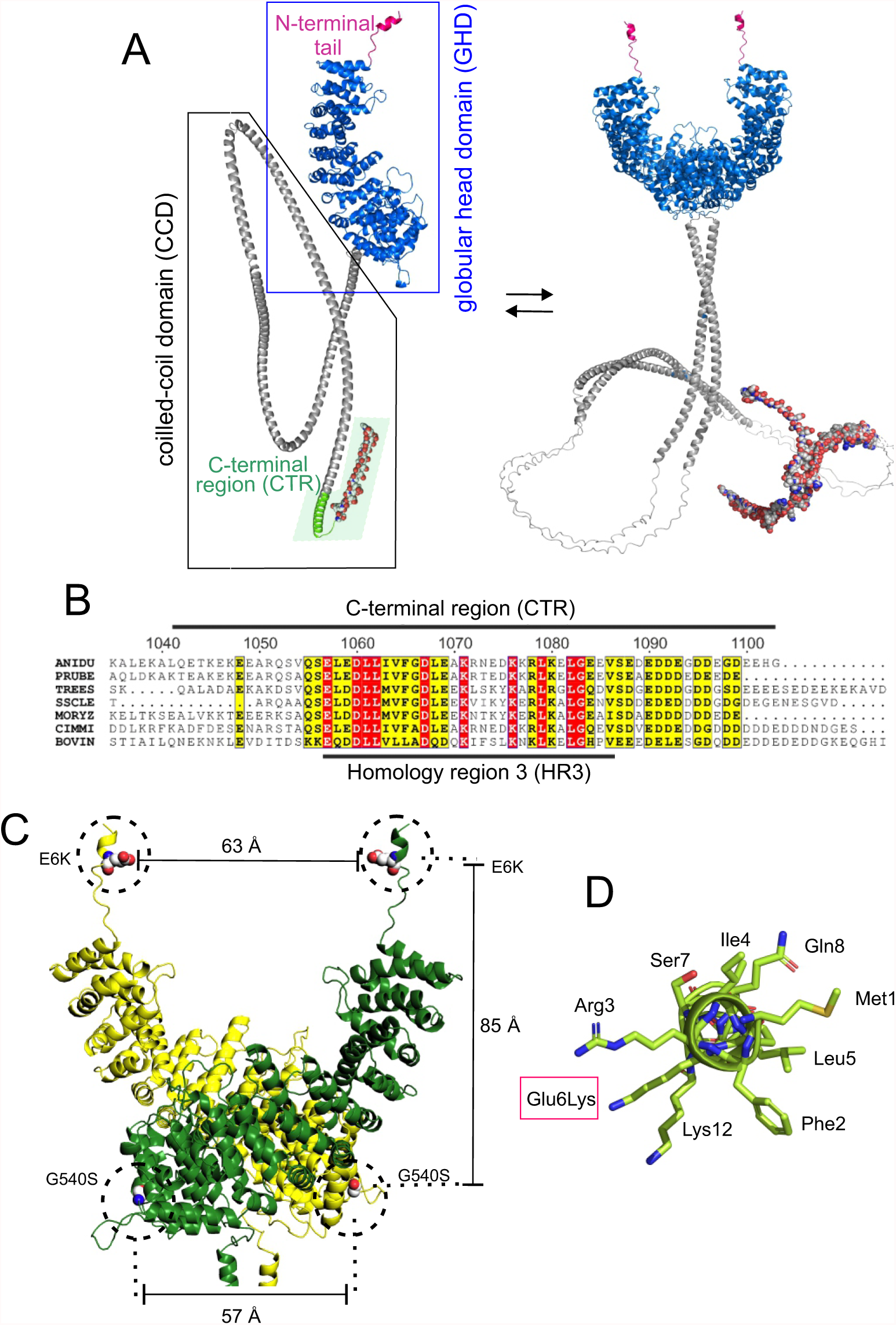
Localization of the amino acid substitutions within the Uso1 AlphaFold2 structure. (A). AlphaFold2 cartoon representations of A. nidulans Uso1 in monomeric and dimeric forms. Confidence estimations for this and other models are detailed under extended data S2. Red, N-terminal tail; marine blue, globular head domain; gray, coiled-coil; green, limit of the CTR. The rest of the CTR is shown as surface representation. (B). Amino acid alignment of fungal sequences with mammalian p115 showing strong conservation within the CTR: ANIDU, *Aspergillus nidulans;* PRUBE, *Penicillium rubens*; TREES, *Thrichoderma ressei*; SSCLE, *Sclerotinia scleriotorum*; MORYZ, *Magnaporthe oryza*e; CIMM, *Coccidioides immitis;* BOVIN, *Bos taurus*. (C). Position of the Gly6Lys and Gly540Ser substitutions. Only the GHD of dimeric full- length Uso1 are shown. The two different chains are colored in green and yellow, respectively. Distances between mutated residues are displayed in armstrongs. (D). The N-terminal amphipathic *α*-helix affected by the Glu6Lys substitution.

Even though the GHD contains Glu6 and Gly540 in the N-terminal helix and at the beginning of armadillo *α*-helix 29, respectively, intramolecular or intermolecular (in the context of a homodimer, see below) distances between these residues are long, arguing against the possibility that they bind a common target as components of the same interaction surface (Figure 2C). Indeed, the synthetic positive effect of the mutations would be consistent with their rescuing viability through different mechanisms. The previously unnoticed short *α*-helix predicted by AlphaFold between Phe2 and Lys12 is amphipathic (Figure 2D). Glu6 lies on the polar side of this helix, such that Glu6Lys increases its overall positive charge (three of the four polar residues are Lys or Arg). As discussed below, it is tempting to speculate that this helix facilitates membrane recruitment.

### Coiled-coil mediated dimerization of Uso1: the globular head is monomeric

It has been proposed that p115 alternates between closed and open conformations to hide or expose a RAB1 binding site present in the CCD (Beard et al., 2005). This would be mediated by intramolecular interactions between the globular domain and the C-terminal acidic region, which would be disrupted by the competitive binding of golgins GM130 and giantin to the latter. We addressed whether E6K/G540S promotes a conformational change in Uso1, or, alternatively, a change in the oligomerization status of the protein, by analytical ultracentrifugation. We designed seven constructs carrying a C-terminal His tag (Figure 3). Two corresponded to the full-length protein with or without E6K and G540S substitutions. The second pair included wild-type and doubly-substituted versions of C-terminally truncated Uso1 lacking the CTR (Uso1ΔCTR and Uso1^E6K/G540S^ΔCTR). The third corresponded to wild-type and doubly-substituted versions of the globular domain, denoted Uso1 GHD and Uso1 GHD^E6K/G540S^. The seventh construct corresponded to the CCD domain (i.e. CC + CTR) and was denoted Uso1 CCD. All seven proteins were expressed in bacteria, purified by nickel-affinity and size-exclusion chromatography and analyzed by sedimentation velocity ultracentrifugation. These experiments revealed that all protein preparations were essentially homogeneous, and thus they were used to determine the corresponding Svedberg coefficients. In addition, by dynamic light scattering we determined the translational diffusion coefficients of the constructs. With these values we deduced the molecular mass of the different proteins using Svedberg’s equation.

**Figure 3:**
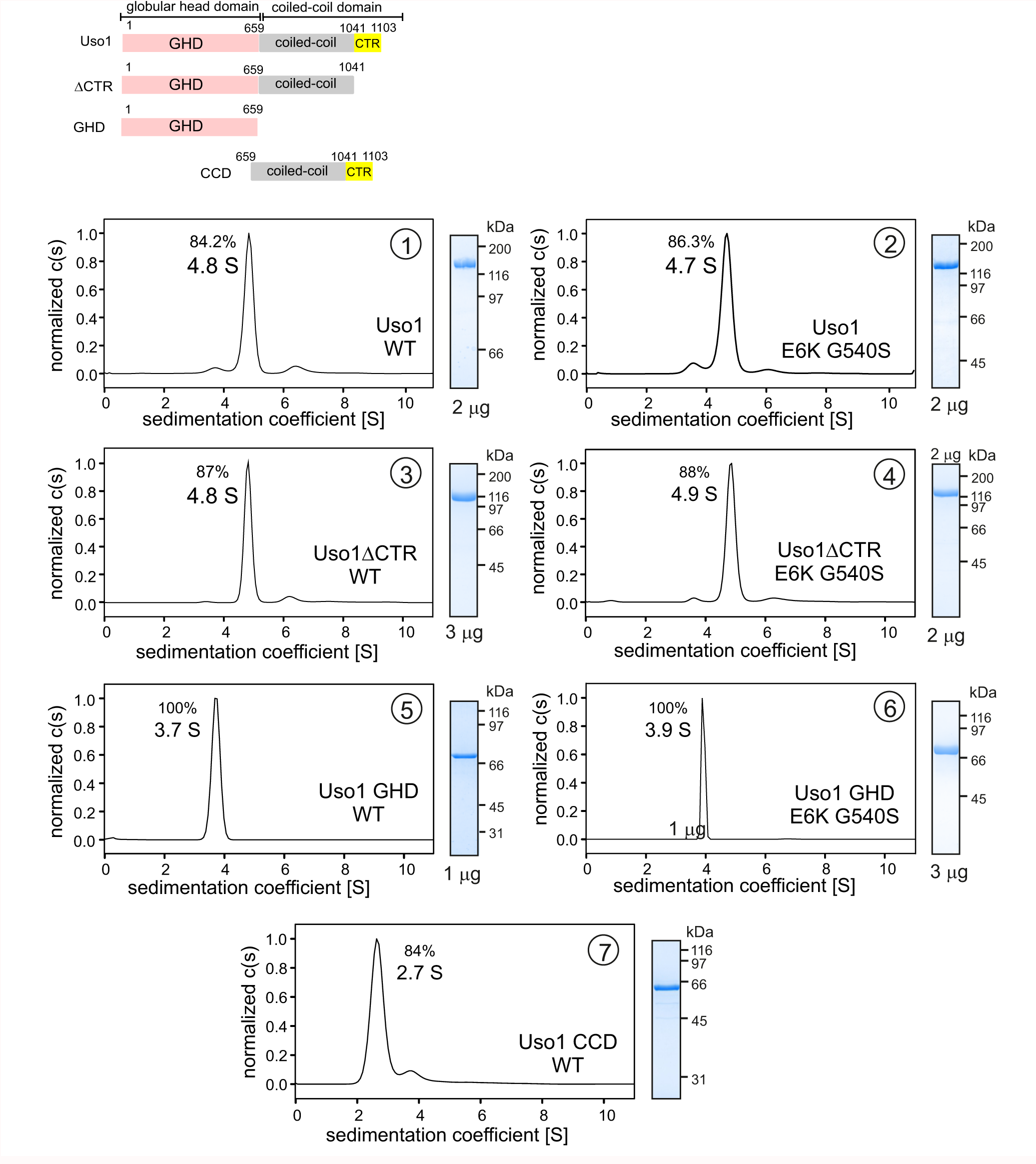
Determining molecular masses and oligomerization status of the different Uso1 constructs by velocity sedimentation analysis. The different panels display the sedimentation profiles of the protein being analyzed, with % of the main species, scheme of the different constructs and their limits and pictures of Coomassie stained-gels showing the purity of the protein preparations. The table below depicts biophysical parameters of the constructs used to obtain relative molecular masses. *sexp* is the experimentally determined Svedberg coefficient; *Dexp*, translational diffusion coefficient of the main species; *Mr*, molecular mass deduced from Svedberg equation; M1 predicted molecular mass of the monomer; n = (*Mr* /M1).

Wild-type and E6K/G540S full-length Uso1s showed the same sedimentation coefficients, demonstrating that the mutations do not induce a large conformational change that would have been reflected in changes in sedimentation velocity due to differences in frictional forces. Molecular masses deduced from the Svedberg equation indicated that these full-length proteins are homodimers, in agreement with previous literature (Figure 3). Ablation of the conserved C-terminal region (CTR) did not result in any significant change in the sedimentation coefficient (Figure 3, panels 3 and 4 vs. 1 and 2), irrespective of the presence or absence of the substitutions, discarding the model in which the CTR would interact with the GHD to maintain a hypothetical closed conformation (Beard et al., 2005). In addition, the molecular masses of the ΔCTR proteins correspond to a dimer, implying that the acidic region is not involved in dimerization either.

Notably the GHD, whether wild-type or mutant, behaved as a monomer (Figure 3, panels 5 and 6), which has important implications described below. In contrast, the coiled-coil domain, with a predicted molecular mass of 52 kDa, behaves as a dimer of *ca.* 100 kDa (Figure 3, panel 7). The sedimentation coefficient of the CCD is markedly slower than that of the 70 kDa monomeric GHD, suggesting an elongated shape. These observations, together with the dimeric nature of the construct lacking the CTR, showed that dimerization is mediated by the CCD. The absence of 443 residues corresponding to the CCD plus CTR domains in the GHD construct and the monomeric nature of the latter compared to full-length Uso1 dimer did not result in a commensurate decrease in sedimentation coefficient, which changed from 4.8 S to 3.7 S in the wild-type (note that the change in *Mr* goes from 246 kDa in full-length Uso1 to only 68 kDa of the GHD)(Figure 3). These data strongly support AlphaFold2 predictions depicting Uso1 as a dimer with a globular head and an extended coiled-coil that would retard sedimentation of the protein very substantially.

As with full-length Uso1, the double substitution did not alter the sedimentation coefficient of the GHD (Figure 3, panels 5 and 6). To buttress the conclusion that the GHD is a monomer irrespective of the presence or absence of the mutations, we performed sedimentation velocity experiments using different protein concentrations ranging from 0.5 to 5 μM (Figure 3—figure supplement 1, A and B). In all cases the GHD behaved as a monomer. Sedimentation profiles of Uso1 GHD lacking the His-tag showed a similar behavior, establishing that the monomeric state of the mutant is not due to the tag at the C-terminal position hindering dimerization (Figure 3—figure supplement 1,C). Therefore, sedimentation experiments did not detect any change in tertiary or quaternary structures between wild-type and mutant GHD, which is important for the interpretation of genetic data that will be discussed below.

In summary, (i) Uso1 is a dimer; (ii) The C-terminal acidic region is dispensable for dimerization and does not mediate an equilibrium between closed and open conformations; (iii) The globular domain of Uso1 is a monomer; (iv) The coiled-coil domain of Uso1 is a dimer; (v) the double E6K G540S substitution does not promote any conformational shift in Uso1, nor does it result in a change in the oligomerization state of the protein.

### The punctate pattern of localization of USO1-GFP is dependent on RAB1

The membranous compartments of the Golgi are not generally stacked in fungi, permitting the resolution of cisternae, which appear as punctate structures in different steps of maturation, by wide-field fluorescence microscopy (Losev et al., 2006; Matsuura-Tokita et al., 2006; Pantazopoulou and Peñalva, 2011; Pinar et al., 2013; Wooding and Pelham, 1998). While Uso1 is predicted to localize to the Golgi, studies of its localization in fungi are limited (Cruz-García et al., 2014; Sánchez-León et al., 2015). Therefore, we tagged the *A. nidulans uso1* gene endogenously with GFP Figure 4A and video 1 depicting a software-shadowed 3D reconstruction of a Uso1-GFP hypha, as well as consecutive sections of deconvolved z-stacks in Figure 4B show that Uso1-GFP localizes to puncta polarized towards the tip, often undergoing short-distance movements (see Figure 4C and Figure 4—figure supplement 1). These puncta are smaller and more abundant than those reported for other markers of the Golgi, which suggested that they might represent domains rather than complete cisternae. Notably, 3D (x, y, t) movies revealed that Uso1 puncta are transient, recurrently appearing and disappearing with time (Figure 4C). That this recurrence did not reflect that the puncta go in-and-out of focus was established with 4D (x, y, z, t) movies, which revealed a similar behavior of Uso1 irrespective of whether 3D or 4D microscopy was used (video 2) Uso1 foci with confidence, we constructed movies with middle planes only (i.e. 3D x, y, t series). After careful adjustment of live imaging conditions, we achieved a 2 fps time resolution with relatively low bleaching for time series consisting of 400 photograms (video 3). These conditions sufficed to track Uso1 puncta over time using kymographs traced across linear ROIs covering the complete width of the hyphae (Figure 4C). However, as the abundance of Uso1 puncta made automated analysis of Uso1 maturation events troublesome, we tracked them manually with the aid of 3D (x,y,t) representations generated with Imaris software combined with direct observation of photograms in movies (Figure 4C and Figure 4—figure supplement 1). The boxed event magnified in Figure 4E (see video 4) illustrates a prototypical example. The right Figure 4E montage shows frames corresponding to this event for comparison. We analyzed *n =* 60 events, which gave an estimation of the average half-life of Uso1 residing in puncta of 60 sec +/- 25.26 S.D. (Figure 4D).

**Figure 4:**
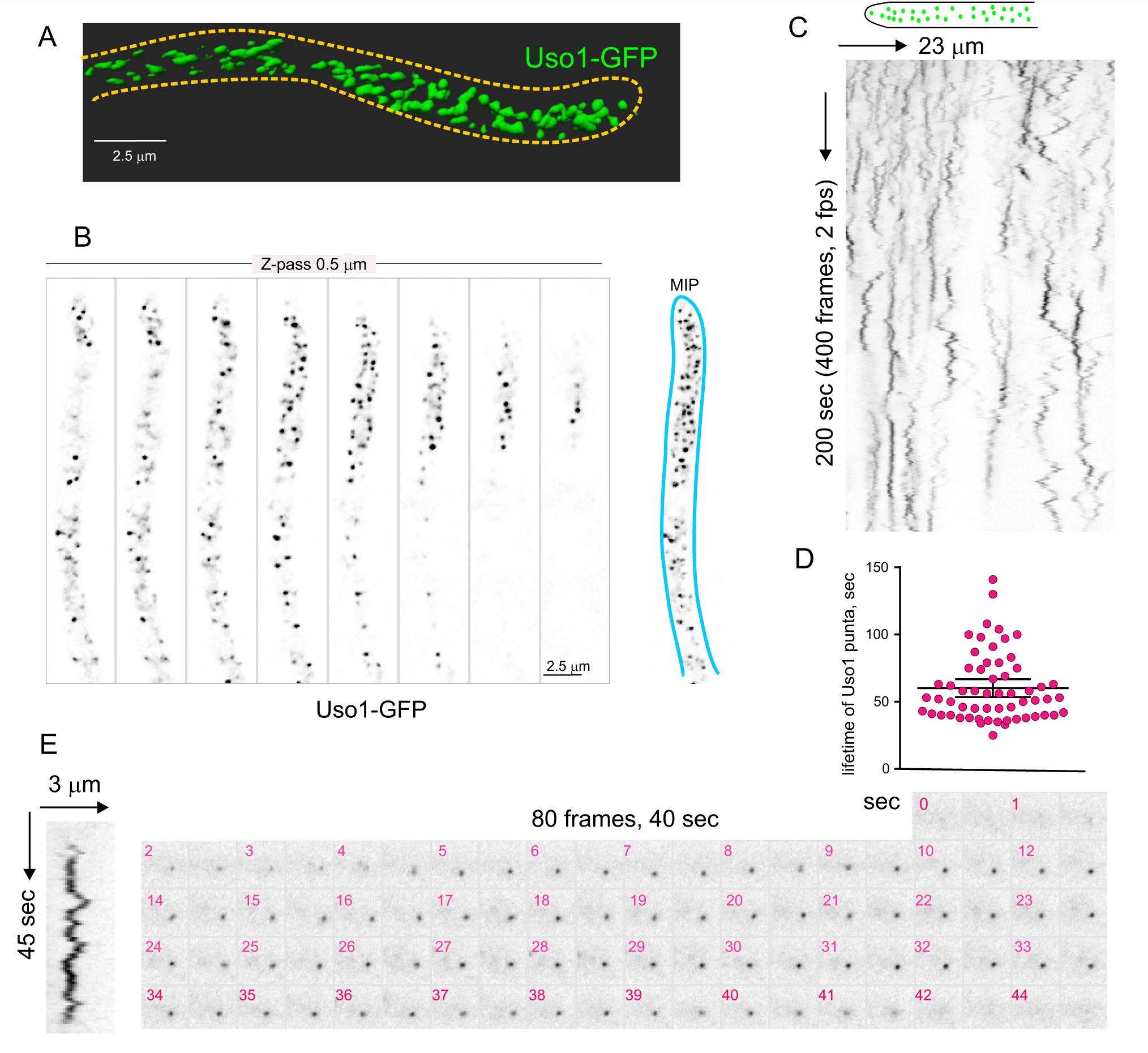
Subcellular localization of Uso1. (A). Uso1-GFP localizing to punctate cytoplasmic structures, 3D shaded by software. (B). Sections of a deconvolved Z-stack and its corresponding MIP. Uso1-GFP in inverted greyscale for clarity (C). Kymograph showing the transient recruitment of Uso1 to punctate cytoplasmic structures. (D). Average time of residence of Uso1 in these structures. Error bars, 95% CI. (E). Example of one such structures visualized with a kymograph and with the corresponding movie frames (Movie 4).

Nakano and co-workers have proposed that the transfer of lipids and proteins between ER exit sites (ERES) and the early Golgi occurs through a kiss-and-run mechanism (Kurokawa et al., 2014). Because Uso1-GFP punctate structures resemble, in size and abundance, ER exit sites labelled with COPII components, we studied Uso1-GFP cells co-expressing Sec13 endogenously labeled with mCherry (Bravo-Plaza et al., 2019). The maximal intensity projection (MIP) shown on Figure 5A, and video 5 show that the two markers are closely associated, but only in a few instances they showed colocalization. These examples did not represent simple overlap, as they were found to colocalize in the Z dimension using orthogonal views or montages (Figure 5B and C). These observations have not been pursued further with time-resolved sequences, but at the very least we can conclude that the reporters are closely associated in space. In view of this, we determined that Uso1 structures originate downstream of COPII-mediated ER exit. Therefore, we investigated, using *sarA*6, a temperature-sensitive allele of the gene encoding *A. nidulans* SAR1 (Hernández-González et al., 2014), whether the punctate Uso1 structures are dependent on this master GTPase regulating COPII biogenesis. Figure 5D shows that this is indeed the case; the number of Uso1-GFP puncta was significantly reduced relative to the wt when cells were shifted from 28°C to 37°C, indicating that Uso1 populates a membrane compartment with Golgi identity.

**Figure 5:**
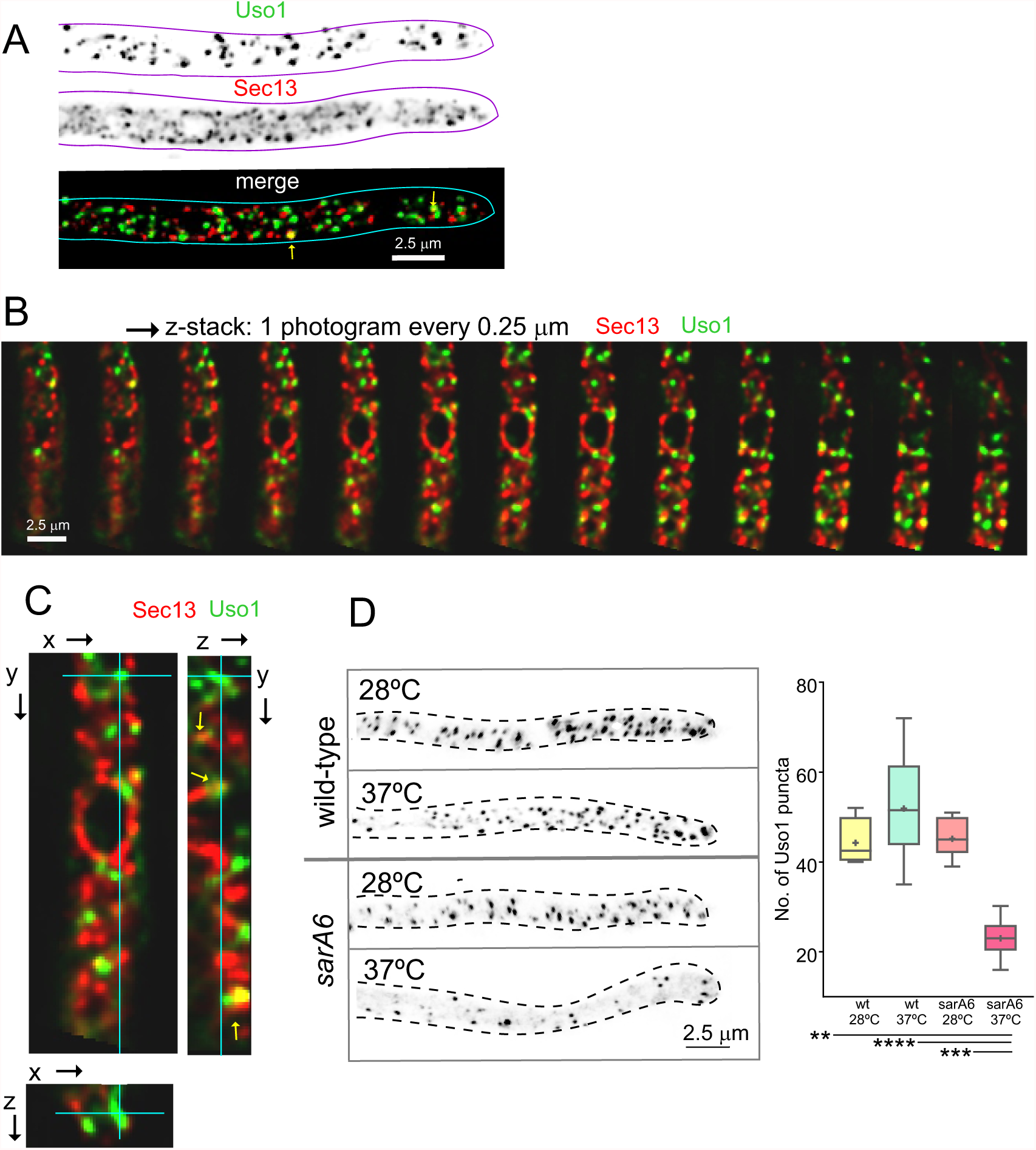
Uso1 puncta do not colocalize with ERESs. (A). Low extent co-localization of Sec13 ERES and Uso1 structures. Z-stacks for the two channels were acquired simultaneously, deconvolved and represented as MIPs. Two rare examples of colocalization are arrowed. (B). Photograms of a dual channel Z-stack with a Sec13-labeled nuclear envelope focused in the middle plane, illustrating that while some puncta show colocalization, the red Sec13 signal and the green Uso1 signal do not usually overlap. (C). A MIP of the same z-stack showing orthogonal views with some overlapping puncta (arrows). (D). A ts mutation in the *sarA* gene encoding the SarA^Sar1^ GTPase governing ER exit markedly reduces the number of Uso1-GFP puncta upon shifting cells to restrictive conditions. Box-and-whisker plots: Statistical comparison was made using one-way ANOVA with Dunn’s test for multiple comparisons. Whiskers are in Tukey’s style: Only significant differences were indicated, using asterisks.

To determine the ‘sub-Golgi’ localization of Uso1 puncta, we filmed Uso1-GFP along with different Golgi markers (Figure 6). Uso1-GFP showed no overlap (Pearson’s coefficient 0.17 ± 0.06 S.D., *n* = 16 cells) with cisternae labeled with mCherry-Sec7, the late Golgi ARF1 GEF that is a prototypic marker of the TGN (Arst et al., 2014; Day et al., 2018; Galindo et al., 2016; Halaby and Fromme, 2018; Losev et al., 2006; McDonold and Fromme, 2014; Pantazopoulou, 2016; Pantazopoulou and Glick, 2019; Richardson et al., 2016; Richardson et al., 2012) (Figure 6; video 6). In contrast, visual observation of cells expressing mCh-Sed5 and Uso1-GFP revealed substantial, yet incomplete, overlap of the reporters (Figure 6), reflected in a Pearson’s coefficient of 0.44 ± 0.04 S.D., *n* = 15 cells (Figure 6). The Qa syntaxin Sed5 drives fusion of COPII vesicles with early Golgi cisternae, with Qb, Qc and R-SNAREs Bet1, Bos1 and Sec22, and mediates intra-Golgi trafficking, with Qb, Qc and R-SNARES Sft1, Gos1 and Ykt6), respectively (Banfield et al., 1995; McNew et al., 2000; Parlati et al., 2002; Pelham, 1999; Wooding and Pelham, 1998). These data suggest that Uso1 localizes to a subset of early Golgi cisternae/membranes containing Sed5. GeaA^Gea1,2^ is the only *A. nidulans* homologue of the *S. cerevisiae* early Golgi ARF1 GEFs Gea1 and Gea2 dwelling at the early Golgi (Arst et al., 2014; Gustafson and Fromme, 2017; Muccini et al., 2022; Pantazopoulou, 2016; Park et al., 2005; Wright et al., 2014). Overlapping of Uso1 with GeaA^Gea1,2^ was more conspicuous than with Sed5 (Figure 6), which was reflected in an increased Pearson’s coefficient to 0.52 ± 0.06 S.D., *n* = 16 cells. Of note, mammalian GeaA (GBF1) and Uso1 (p115) interact (Garcia-Mata and Sztul, 2003). Taken together, these data indicate that Uso1 localizes to Golgi cisternae in early stages of maturation.

**Figure 6:**
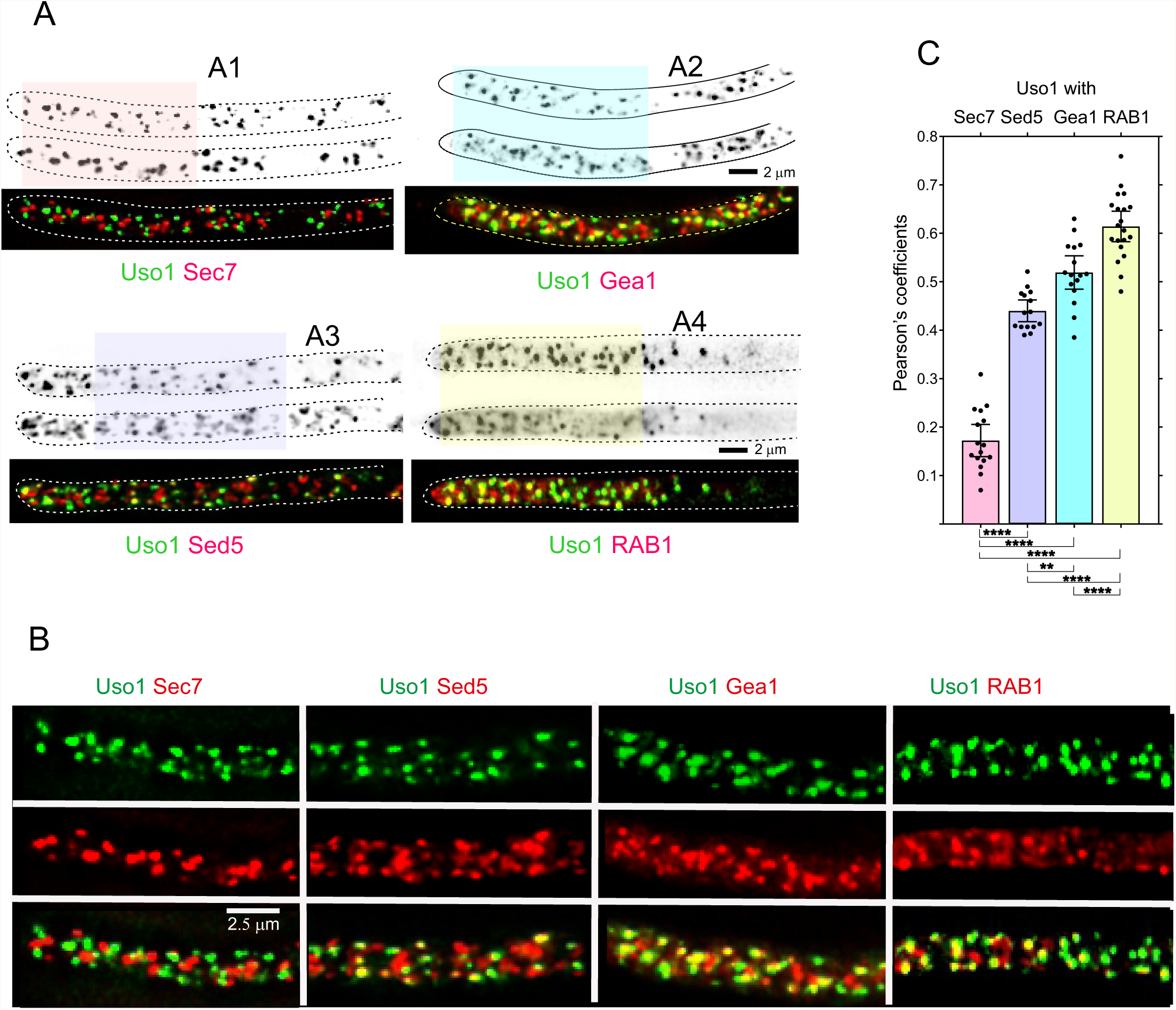
Uso1 localizes to RAB1-containing Golgi cisternae. (A). Tip cells showing Uso1 colocalization with the indicated subcellular markers. Images are MIPs of deconvolved Z-stacks. (B). Magnified images of the color-coded shaded regions of the cells shown in A. (C). Pearson’s coefficients of the different combinations.

In both mammalian cells and in yeasts Uso1/p115 has been shown to be recruited to early Golgi membranes by RAB1, which is activated by the TRAPPIII GEF on COPII vesicles after they bud from the ER (Allan et al., 2000; Cai et al., 2007; Lord et al., 2011; Yuan et al., 2017). Therefore, we imaged Uso1 and RAB1, which revealed that indeed Uso1 colocalized with RAB1 (Pearson’s 0.61 ± 0.07 S.D., *n* = 20 cells) (Figure 6). Altogether, the above microscopy data strongly suggest that Uso1 is transiently recruited to early Golgi membrane domains enriched in RAB1, agreeing with the accepted view that RAB1 acts by recruiting Uso1.

### Uso1 delocalization after RAB1 impairment rescued by E6K G540S

We next tested if the subcellular localization of Uso1 is dependent on RAB1, and if this dependency can be bypassed by E6K/G540S. To this end we first showed that in a *RAB1^+^* background endogenously tagged wild-type and E6K/G540S Uso1-GFP have the same punctate localization pattern (Figure 7A), and that both supported vigorous wt growth (Figure 7B lanes 1, 3, 5 and 7), indicating that the tagged proteins are functional. Next, we introduced in these strains *rab1^ts^* (Jedd et al., 1995; Pinar et al., 2013) by crossing. This allele, which completely prevents growth at 37°C, is a hypomorph at permissive (25-30°C) temperatures, which permitted testing RAB1 dependence under standard microscopy conditions (28°C). Figure 7A shows that wt Uso1-GFP was largely delocalized to the cytosol by *rab1^ts^*. This establishes that Uso1 localization to membranes is subordinated to RAB1. Notably, the wt punctate pattern of Uso1 localization was restored by the Uso1 E6K/G540S mutant substitution, correlating with correction of the synthetic growth defect (Figure 7B, compare lanes 2, 4, 6 and 8). All these data indicated that the localization of Uso1 is compromised when RAB1 function is impaired, and that the E6K/G540S substitutions augment Uso1 affinity for a membrane anchor(s) independent of RAB1. They additionally suggest that the principal physiological role of RAB1 is ensuring the proper localization of Uso1.

**Figure 7:**
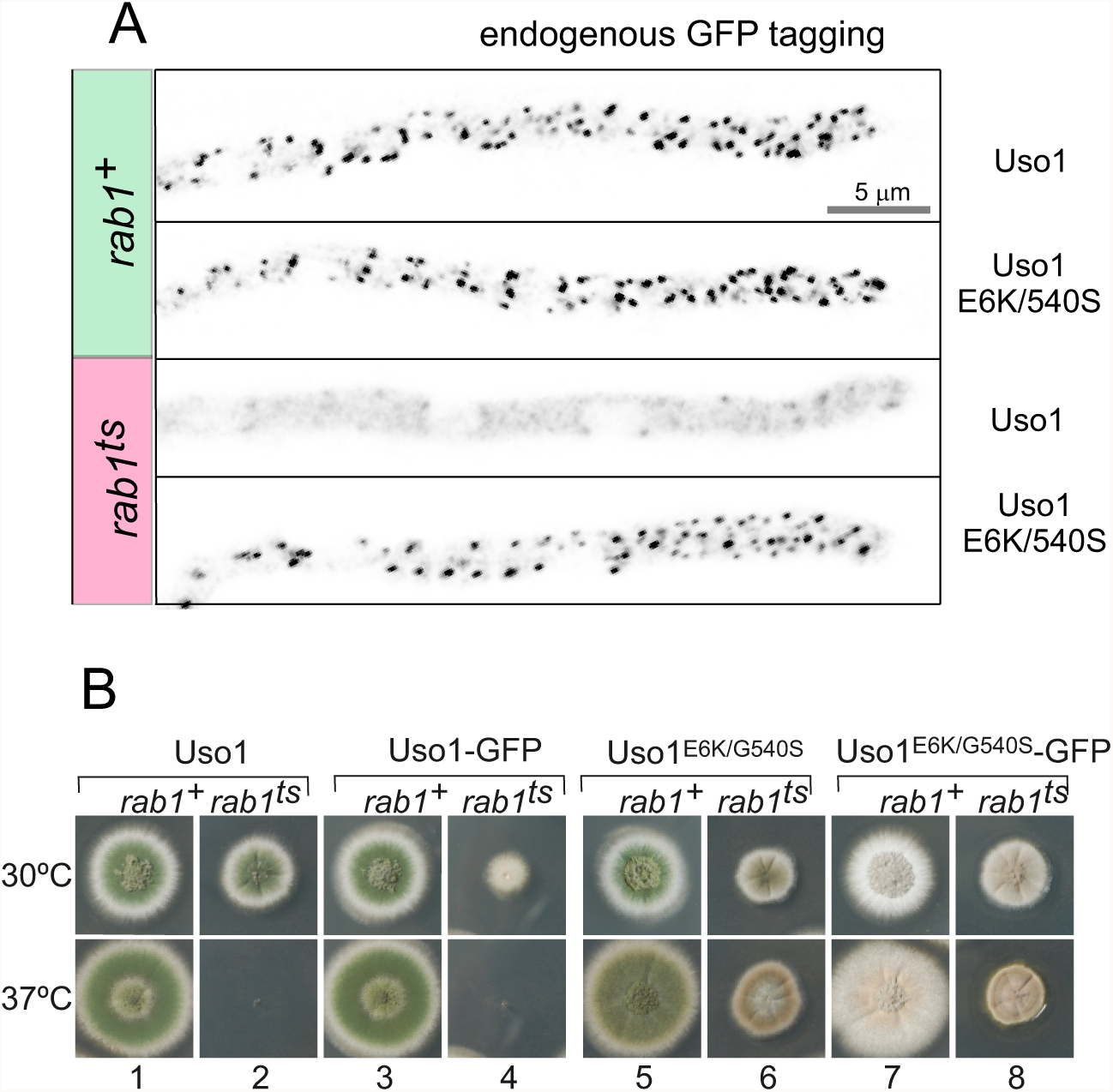
Uso1 localization to punctate structures is dependent on RAB1. (A). Complete de-localization of Uso1-GFP to the cytosol by rab1ts and relocalization by E6K/G540S. (B). Uso1-GFP and *rab1ts* show a synthetic negative interaction that is rescued by the E6K/G540S double substitution. Strains in lanes 7 and 8 carry the *wA2* mutation resulting in white conidiospores.

Delocalization of Uso1 in the *rab1^ts^* background was not solely dependent on RAB1. Wild-type *uso1-GFP* displayed a synthetic negative interaction with *rab1^ts^* (Figure 7B, compare lanes 2 and 4 at 30°C), suggesting that the presence of GFP in the C-terminus interferes with a RAB1-independent mechanism that facilitates its recruitment to membranes (see below)

### Genetic evidence that a network involving the CTR and the Grh1/Bug1 golgin contributes to the recruitment of Uso1 to membranes

To follow up the above observation, we focused on golgins. In mammalian cells, the C-terminal region of p115 interacts with GM130, a golgin which is recruited to the early Golgi by GRASP65 (Beard et al., 2005). The equivalent proteins in budding yeast are denoted Bug1 and Grh1 (Behnia et al., 2007)(Figure 8A).

**Figure 8.**
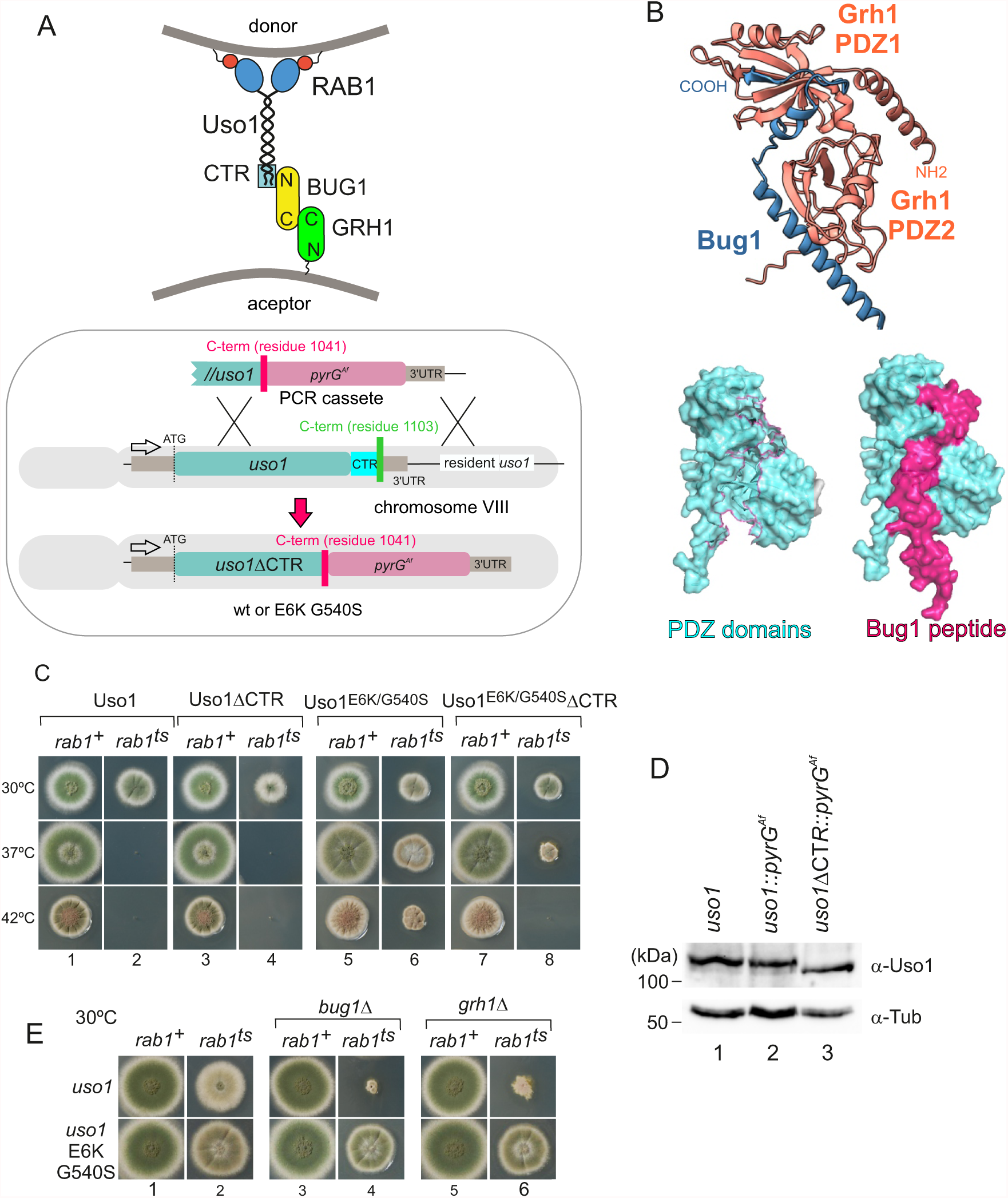
Genetic evidence showing that the CTR region of Uso1 contributes to its recruitment to membranes. (A). Top, scheme of the predicted interactions. Bottom, engineering a gene-replaced allele lacking the CTR domain by homologous recombination. (B). The Bug1 C-terminal residues fit into the groove formed between the two Grh1 PDZ domains and into the pocket of the N-terminal PDZ domain (PDZ1). (C). A gene-replaced *uso1ΔCTR* allele encoding a protein truncated for the CTR domain shows a synthetic negative interaction with *rab1ts*. (D). Western blot analysis. Removal of the CTR does not result in Uso1 instability. (E). *bug1*Δ and *grh1*Δ show a synthetic negative interaction with *rab1ts* that is rescued by the double E6K/G540S substitution in Uso1

GRASP65 contains two C-terminal PDZ [post synaptic density protein (PSD95), *Drosophila* disc large tumor suppressor (Dlg1), and zonula occludens-1 protein (zo-1)] domains, which bind an also C-terminal peptide in GM130 (Hu et al., 2015).. Similar to its metazoan counterparts, *Aspergillus* Grh1 contains an N-terminal *α*-helix and two PDZ domains, in this case followed by ∼130 disordered residues (Figure 8B, Figure 8—figure supplement 1A and C). We modelled the Grh1-Bug1 interaction using AlphaFold2. Residues 666-675 of the C-terminal peptide of Bug1 bind to a hydrophobic cleft located between PDZ1 and PDZ2. Bug1 Leu668 and 670 coordinate their side chains with residues from both PDZ domains (e.g. Phe45 and Trp44 in PDZ1 and Trp171 and Val179 in PDZ2). A second interaction involves Bug1 C-terminal residues 683-690 fitting within a second groove in PDZ1, such that the four C-terminal residues form a β-strand extending the β-sheet of N-terminal PDZ1 domain (Figure 8B, Figure 8—figure supplement 1B). Further genetic evidence that a network of interactions similar to that acting in yeast and mammalian cells operates in the *A. nidulans* ER-Golgi interface was obtained by constructing strains with combinations of gene-replaced alleles. These consisted of *uso1ΔCTR* encoding Uso1 lacking the C-terminal region (residues 1-1041) and containing or not E6K G540S, *rab1^ts^*, and deletion alleles of the *Aspergillus BUG1* (AN7680) and *GRH1* (AN11248) genes (Figure 8A). Combining *rab1^ts^* with *uso1ΔCTR* resulted in a synthetic negative interaction at 30°C akin to that seen with *uso1-GFP* (lanes 2 and 4 in Figure 8C). That the E6K/G540S double substitution rescued this negative interaction strongly indicates that the CTR cooperates with RAB1 in the recruitment of Uso1 to membranes (Figure 8C, lanes 2, 4, 6 and 8). We note that the control wild-type strain used in these experiments contains a construct completely analogous to the mutant allele, ruling out that the genetic manipulation (for example the introduction, linked to the *uso1* locus, of a selection marker, potentially chromatin- disruptive) is causative of the observed phenotype. Another trivial explanation that we ruled out by Western-blot analysis was that the deletion off the 62 C-terminal residues in *uso1ΔCTR* resulted in increased degradation, which it did not (Figure 8D).

If interactions involving the CTR of p115 were conserved in fungi, *BUG1* (GM130 equivalent) and *GRH1* (GRASP65) should also show a synthetic negative interaction with *rab1^ts^*. Figure 8E (lanes 1,3 and 5) shows that neither *bug1*Δ nor *grh1*Δ affects growth. However, both deletion alleles showed a strong synthetic negative interaction with *rab1^ts^* (Figure 8E, lanes 1,3 and 5). Remarkably, the synthetic negative phenotype was rescued by the presence of E6K/G540S substitutions in Uso1, further suggesting that they promote Uso1 recruitment to its locale of action, compensating for the loss of the Bug1-Uso1 CTR interaction. We conclude that interaction involving the CTR of Uso1 and the BUG1/GRH1 complex cooperates with RAB1-mediated mechanisms to recruit Uso1 to membranes.

### The globular head domain (GHD) of Uso1 carrying the double E6K/G540S substitution supports cell viability

Uso1 has traditionally been considered the archetype of a coiled-coil tether recruiting ER-derived vesicles to the *cis-*Golgi. In *S. cerevisiae*, *uso1-1* (Sapperstein et al., 1996), is a C-terminally truncating, conditionally-lethal *ts* allele whose encoded protein still retains 20% of the coiled-coil region. In contrast, *uso1-12* and *uso1-13* removing the complete coiled-coil region are lethal (Seog et al., 1994). Thus, we asked whether the double E6K/540S substitution bypasses the requirement of the coiled-coil region. To this end, we constructed, by gene replacement, a *uso1^GHD^* allele expressing a protein truncated immediately after the GHD, lacking residues 660-1103. By heterokaryon rescue, we demonstrated that this allele is lethal (Figure 9A). Unexpectedly, given that the *rab1^ts^* suppressor substitutions lie outside the coiled-coil region, the equivalent allele containing E6K/G540S sufficed for the fungus to survive at 30°C. This result has two key implications: that the coiled-coil of Uso1 is not essential for survival and as the GHD of Uso1 is a monomer in solution, it follows that dimerization of Uso1 is not essential either (see also below).

**Figure 9:**
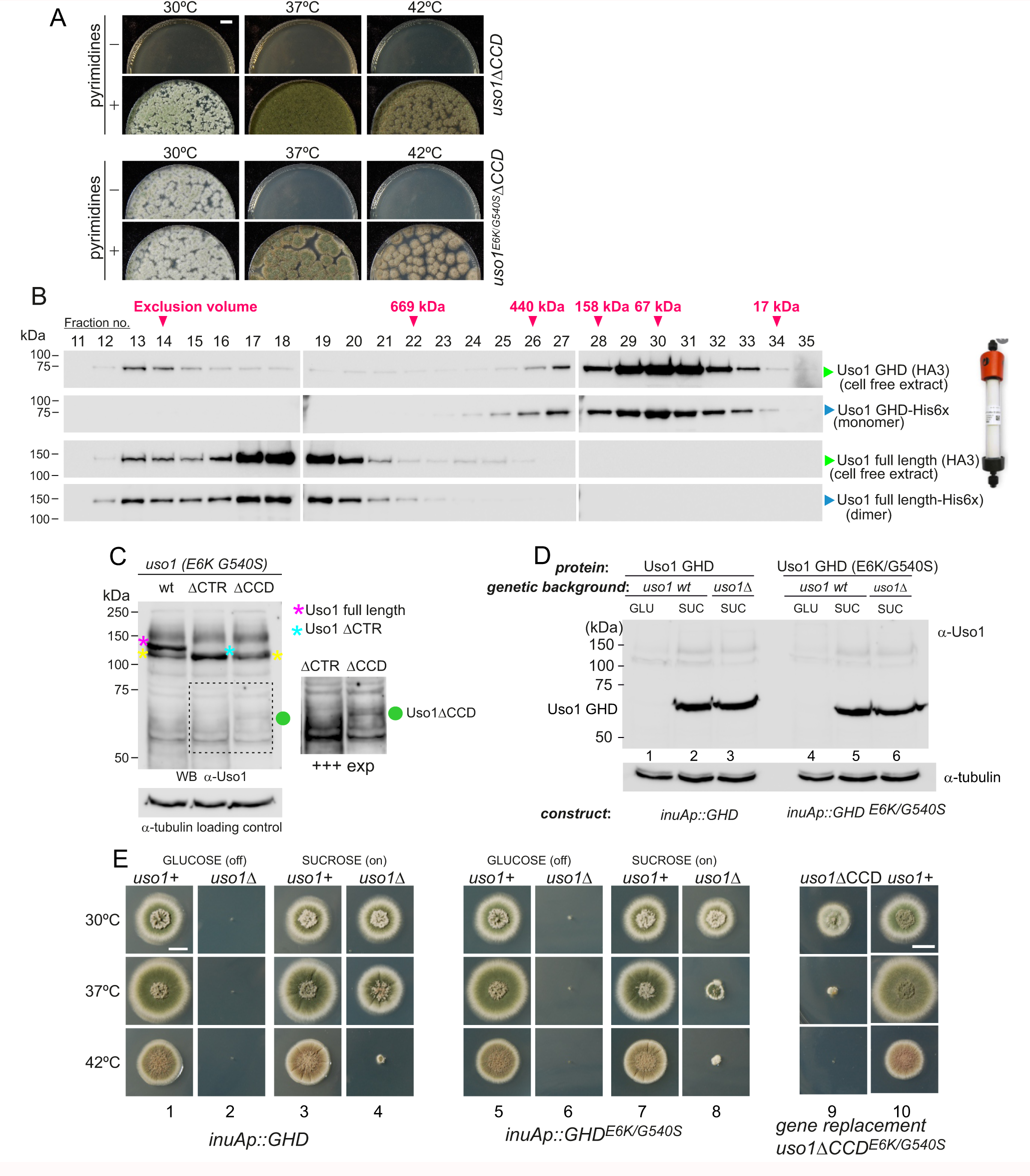
The GHD of Uso1 is sufficient to support cell viability. (A). Gene-replaced *uso1^GHD^* allele carrying the double E6K/G540S substitution is sufficient to rescue viability at 30°C, but not at higher temperatures. (B). The Uso1 GHD is a monomer in vivo. Fractions collected from Superose columns loaded with the indicated protein extracts and reference His-tagged proteins were collected and analyzed by western blotting with *α*-HA and *α*-His antibodies. (C) Truncating Uso1 after the GHD results in markedly reduced protein levels, as determined by *α*-Uso1 GHD western blotting. The band (yellow asterisks) moving slower than Uso1 (magenta asterisk) and at nearly the same position of Uso1 ΔCTR (blue asterisk) represents cross- reacting contaminants unrelated to Uso1. The right panel shows a longer exposure for the indicated region, to reveal the faint GHD band (green dot). (D). Overexpression of Uso1, wild-type and E6K/G540S mutant, under the control of the *inuA* promoter, which is turned off on glucose and induced on sucrose. Western blots reacted with *α*-Uso1 GHD antiserum. (E). Overexpressed GHD, be it E6K/G540S or wild-type, as the only source of Uso1 supports viability.

In view of these unexpected results we wondered whether the structure of the GHD synthesized in bacteria differed from the physiological form in *Aspergillus*, such that the GHD were a dimer *in vivo.* To address this possibility, we ran an *Aspergillus* cell-free extract expressing HA3-tagged Uso1 GHD through a Sepharose column. As control, we ran in parallel a sample of bacterially-expressed His-tagged GHD used in sedimentation velocity experiments. Western blot analysis of the fractions (Figure 9B) demonstrated that both proteins eluted at the same position, corresponding to that expected for a globular protein with the size of the GHD. As controls we ran in the same column a similar pair of proteins corresponding to full length Uso1. The bacterially-expressed and the native Uso1 proteins also co-eluted (Figure 9B), but this time at a position corresponding to a highly elongated dimer, consistent with sedimentation velocity experiments. Thus, *in vivo*, the GHD, expressed from its own promoter, is a monomer, and it is sufficient to sustain viability if it carries the double mutant substitution that bypasses the requirement for RAB1. These data strongly argue against tethering being *the* essential physiological role that Uso1 plays.

We next investigated why the E6K/G540S GHD suffices for viability only at 30°C. Often thermo-sensitivity results from protein instability, which is enhanced at high temperatures. Western blot analysis of the allele-replaced strain expressing E6K G540S GHD as the only source of Uso1 revealed that levels of the truncated Uso1 mutant were minuscule relative to the wt or to the equivalent *ΔCTR* allele (Figure 9C). We reasoned that increasing expression would result in E6K/G540S GHD supporting growth over a wider range of temperatures. Thus, we drove its expression with the promoter of the inulinase *inuA* gene, which is inducible by the presence of sucrose in the medium and almost completely shut off on glucose (Hernández-González et al., 2018; Peñalva et al., 2020). Initially we tested wild-type and E6K/G540S GHD in a *uso1+* background. This had no phenotypic consequences despite the fact that western blots confirmed that that the truncated proteins were being overexpressed (Figure 9D; Figure 9E, lanes 1, 3, 5, 7 and 10;). Then we proceeded to delete the resident *USO1* gene in the wild-type and mutant GHD overexpressing strains. As expected, neither of the resulting pair of strains was able to grow on medium with glucose as the only carbon source (Figure 9E, lanes 2 and 6). Notably, the strain expressing E6K/G540S GHD as sole Uso1 source grew essentially as the wild-type and 30°C and, although debilitated, was viable at 37°C, showing a substantial improvement of the growth capacity displayed by the gene-replaced mutant (Figure 9, lanes 7,8 and 9). Thus, if expressed at sufficiently high levels, E6K/G540S GHD maintains viability at the optimal growth temperature.

Unexpectedly, the wild-type GHD also rescued the viability of the *uso1Δ* mutant when this was cultured with sucrose as carbon source at 30°C and 37°C. In fact, at 30°C, the *uso1Δ inuAp::GHD* strain grew like the wt (Figure 9E, lanes 3 and 4)(note that these experiments were carried out in a *RAB1+* background), suggesting that increased binding to a Golgi receptor facilitated by mass action compensated for the loss of the coiled-coil region and associated dimerization. The E6K/G540S GHD would have gained affinity for this receptor, explaining why the doubly substituted GHD suppressed mis- localization of Uso1-GFP when RAB1 is compromised, even when its steady-state levels were very low. Forced expression, combined with a potentially increased binding affinity of E6K/G540S GHD to such a hypothetical receptor might be toxic.

### Uso1 is an associate of the early Golgi SNARE machinery, with the double substitution E6K/G540S increasing this association

What is the nature of this hypothetical receptor? To address this question, we screened for interactors of Uso1 among proteins acting at the same functional level (consumption of COPII vesicles by the early Golgi) using a modified version of the S-tag co-precipitation approach that we used to characterize of TRAPP complexes (Pinar et al., 2019)(Figure 10A). We constructed strains expressing wild-type or mutant Uso1, tagged endogenously with the S-tag and, as negative unrelated control, BapH (an effector of RAB11 acting in late steps of the secretory pathway (Pinar and Peñalva, 2017). Then, derivatives of these three strains co-expressing each of the candidate Uso1 GHD targets, tagged with HA3 (also endogenously), were constructed. The resulting panel (Figure 10B) was screened for HA-tagged proteins co-precipitating more efficiently with the E6K/G540S version of Uso1 than with the wild-type, and satisfying the criterium of not co-purifying with BapH. To this end cell-free extracts of these strains were incubated with S-agarose beads that were recovered by centrifugation. Proteins associating with the S-baits were revealed by anti-HA western blotting.

**Figure 10:**
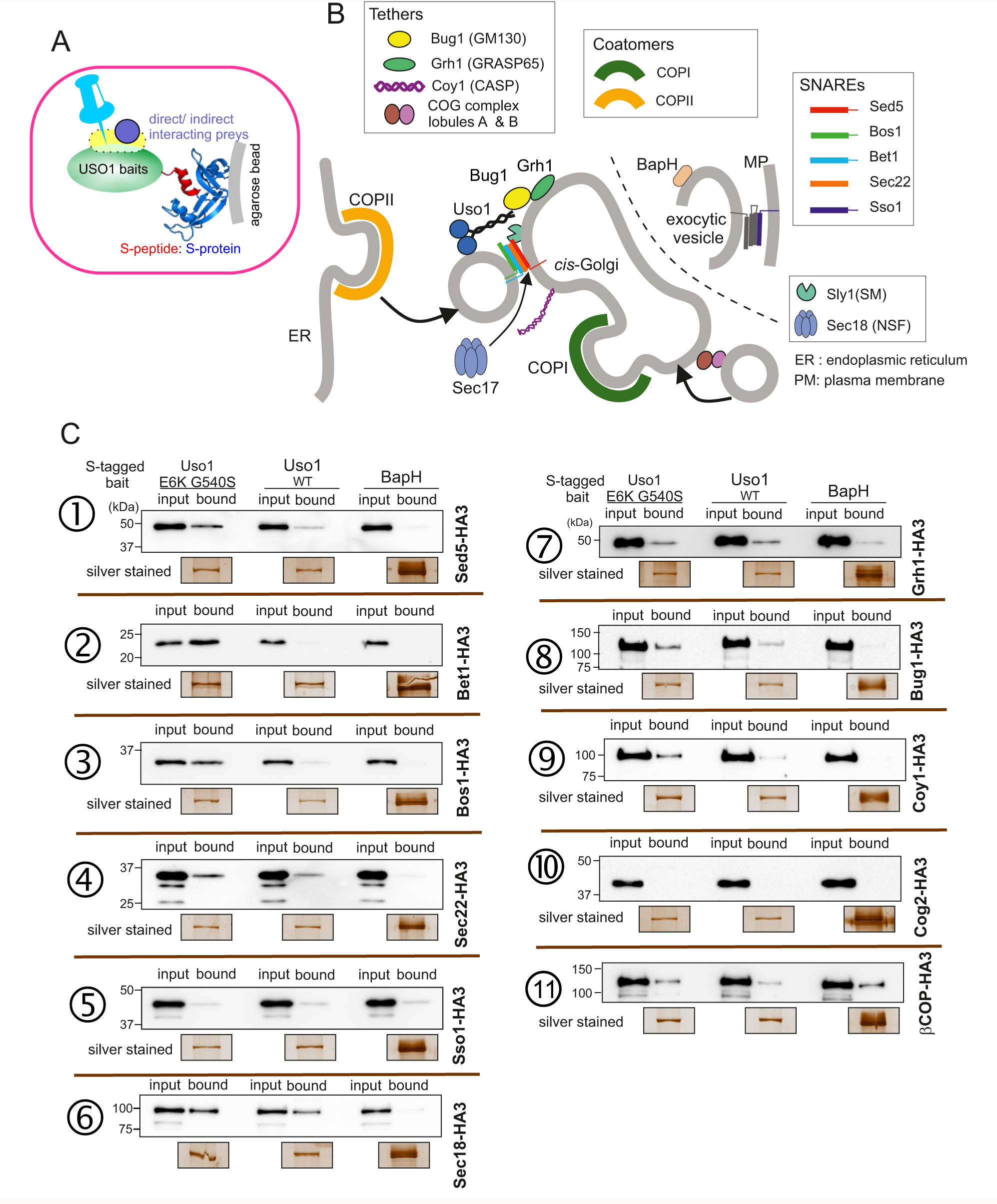
Screening the preferential association of proteins acting in the ER/Golgi interface with E6K/G540S Uso1. (A). S-tagged baits (Uso1, wt and E6K/G540S, and the unrelated protein BapH), expressed after gene replacement, were captured with their associated polypeptides on S-protein agarose beads. Candidate associates, also expressed after gene replacement, were tagged with HA3. (B). Schematic depiction of the proteins listed in these experiments showing their sites of action.(C). Anti-HA3 western blot analysis of the indicated S-bait and HA3-prey combinations. Equal loading of Uso1 proteins was confirmed by silver staining of precipitates. Note that BapH, chosen as negative control, is expressed at much higher levels than Uso1 proteins. Each panel is a representative experiment of three experimental replicates.

That not every protein specifically co-purified with Uso1 baits was demonstrated by the results obtained with the COG component COG2, which did not associate with any of the three baits (Figure 10C, 10). In contrast, *β*-COP was a promiscuous non-specific interactor pulled down by all three baits (Figure 10C, 11). Notably, the screen identified the Golgi syntaxin Sed5 within the specific Uso1 associates (Figure 10C, 1), an association reported previously by others for both p115 and fungal Uso1 (Allan et al., 2000; Sapperstein et al., 1996). That the PM SNARE Sso1 did not interact at all with any of the S-baits demonstrated that Uso1 does not bind promiscuously to syntaxins (Figure 10C, 5). Importantly, Sed5 was brought down more efficiently by E6K/G540S Uso1, and not at all by BapH, even though levels of this unrelated bait, as assessed by silver staining of pull-downs, were markedly higher than those of either Uso1 version. Therefore, Sed5 (or its associates) might represent a potential anchor bound by E6K/G540S Uso1 with increased affinity to compensate for the lack of RAB1-mediated recruitment.

**Figure 11:**
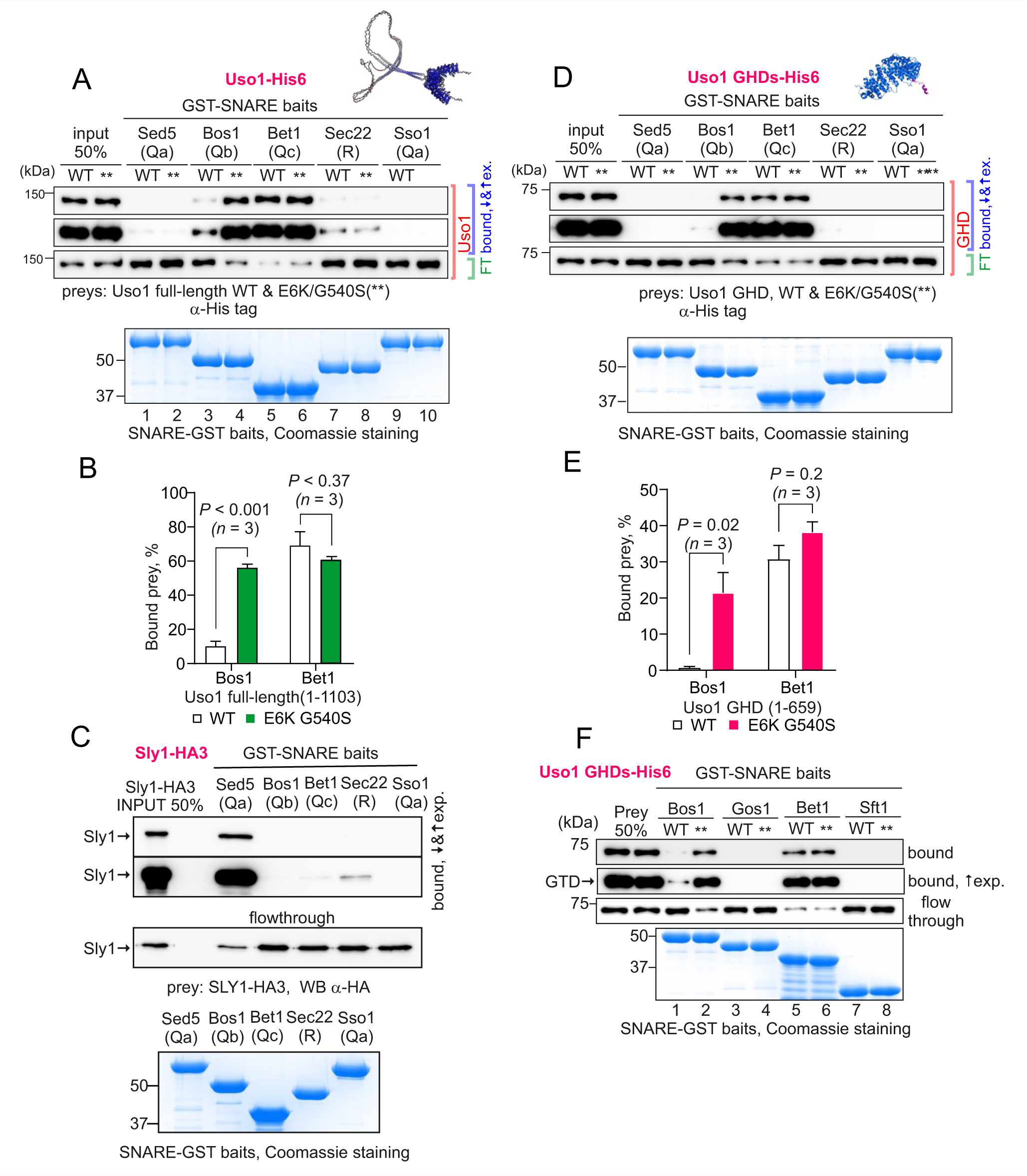
The Uso1 GHD interacts directly with Bos1 and Bet1 SNAREs acting in the ER/Golgi interface. (A). Purified fusion proteins in which the cytosolic domains of the indicated SNAREs have been fused to GST were used in pulldown experiments with His-tagged, purified wild-type and E6K/G540S Uso1. The plasma membrane Qa syntaxin Sso1 was used as negative control. Pulled-down material was analyzed by anti-His western blotting. (B). Quantitation of the above experiment; significance was determined by unpaired *t-*student tests. Error bars represent S.E.M. (C). As in A, but using in vitro synthesized, HA3-tagged Sly1 as prey. Samples were analyzed by anti-HA western blotting. (D). As in A, but using wild-type and mutant GHD as preys, rather than full-length Uso1. (E). Quantitation of the experiment in D. (F). GST pull-down experiment comparing the ability of the GHD to interact with the early Golgi Qb and Qc SNAREs (Bos1 and Bet1), with that of their medial Golgi counterparts (Qb Gos1 and Qc Sft1).

This analysis was extended to other members of the SNARE bundle forming in the ER/Golgi interface with the Qa Sed5: the Qb Bos1, the Qc Bet1 and the R-SNARE Sec22 (Parlati et al., 2002; Pelham, 1999; Tsui et al., 2001). Sec22 was slightly enriched in the E6K/G540S pull-down relative to wild-type Uso1 (Figure 10C, 4). The results with Bos1 and Bet1 were most noteworthy (Figure 10C2 and 3). Both were markedly increased in the E6K/G540S pull-downs, with Bet1 increased most. The AAA ATPase Sec18 disassembling *cis-*SNARE complexes also bound Uso1 and was slightly enriched in the sample of E6K/G540S associates, as was the Uso1-interacting Golgin Bug1, but not its membrane anchor Grh1 (Figure 10, C6-8). We conclude that Uso1 is a component of the SNARE machinery and that this association is augmented by E6K/G540S.

S-tag-coprecipitations in Figure 10 clearly singled out Bet1 (Qc) and Bos1(Qb) as the preys that were most strongly enriched with the mutant E6K/G540S bait relative to wild-type, and therefore with the highest probability of being direct interactors bound with greater affinity by the doubly substituted Uso1 mutant.

During these analyses we also addressed the physiological role of golgins, which were expected to be dispensable for growth, given their functional redundancy (Gillingham, 2018; Gillingham and Munro, 2016; Muschalik and Munro, 2018). We examined the role of three golgins acting at the early Golgi: The Grh1 (AN11248)/Bug1(AN7680) complex, discussed above, Coy1 (the product of AN0762) and Rud3 (AN10186). Consistent with their roles, AlphaFold2 predicts that they form long coiled-coils carrying C-terminal membrane anchors, characteristics of golgins. Coy1 contains a C-terminal TMD anchor and an adjacent CLASP domain that, according to AlphaFold2, consists of *α*-helices (Figure 10—figure supplement 1). Rud3 is a dimer consisting of a long coiled-coil with a ARF1-binding GRIP domain composed of four short helices, near its C-terminus (Figure 10—figure supplemental 1). Ablation of *grh1*, *bug1*, *coy1* or *rud3* did not prevent growth of the corresponding mutants, with only *coy1*Δ displaying a minor growth phenotype (Figure 10—figure supplement 2). Next, we deleted the corresponding genes in a *rab1Δ*

*uso1^E6K G540S^* and tested whether any of the golgins was required for viability rescue. Grh1 and Bug1 were partially required, as their absence precluded rescue at 42° (Figure 10— figure supplement 2). In contrast, RUD3 was not required. With regard to COY1, the fact that combining *coy1*Δ and *rab1Δ uso1^E6K G540S^* results in lethality at 30°C, 37°C and 42°C precluded conclusions on the role of COY1 in suppression. However, we can conclude that proper assembly of *A. nidulans* Golgi cisternae is supported by a redundant set of tethers (Behnia et al., 2007), of which in Coy1 appears to be the least redundant of those tested. Coy1 has been implicated in retrograde traffic within the Golgi itself (Anderson et al., 2017)

### Golgi SNAREs bind directly to the Uso1 GHD; effects of Uso1 E6K/G540S

Work by others implicated Uso1 in the assembly of the early Golgi SNARE bundle (Sapperstein et al., 1996). Thus, prompted by co-association experiments, we predicted that Uso1 would bind directly Bet1, Bos1 and perhaps other SNAREs implicated in the biogenesis of the early Golgi. We anticipated that binding to Bet1 and Bos1 would be insufficient to recruit Uso1 to membranes in the absence of RAB1, but that once reinforced by E6K/G540S, Uso1 would not require RAB1 for its recruitment. To test this possibility, we searched for direct and E6K/G540S-enhanced interactions between the GHD and SNAREs with pull down assays carried out with purified SNARE-GST fusion proteins as baits and Uso1-His6 constructs as preys.

Full-length Uso1 bound, weakly, to the Sec22 R-SNARE and to the Qb Bos1, and very efficiently [with *circa* 70% of the prey being pulled down (Figure 11A, B)] to the Qc Bet1. In contrast, Uso1 did not bind the Qa syntaxins tested, Sed5 and Sso1 (Figure 11A). The absence of interaction between Sed5 and Uso1, be it the wild-type or the E6K/G540S mutant version, strongly indicated that the association detected with S-tag pull-downs between Uso1 and Sed5 is bridged by other protein(s). This absence of binding cannot be attributed to Sed5-GST being incompetent for binding because Sed5-GST was competent in pulling-down highly efficiently its cognate SM protein Sly1, an interaction that did not occur with Sso1-GST (Figure 11C). Notably, the presence of the E6K/G540S double substitution in Uso1 (indicated with ** for simplicity on Fig 11) increased five times the amount of protein retained by the Qb Bos1 bait (Figure 11A and B), whereas interaction with Bet1 did not change (Figure 11A and B) . The double substitution in Uso1 did not promote interaction with Sed5 either. Thus, under normal circumstances Uso1 is able to bind directly to three of the four SNAREs in the ER/Golgi interface, with binding to Bet1 being the strongest. The double E6K/G540S substitution increases binding to Bos1 very markedly and specifically, bringing it to up to the levels of Bet1 without affecting, for example, binding to Bet1 or Sec22. Consistently, the GHD is sufficient to mediate interaction with Bet1 and Bos1, as well as, if E6K G540S-substituted, the increased binding of Bos1 to Uso1 (Figure 11D and E). The GHD did not interact with Sec22, suggesting either that this R-SNARE is recruited by other parts of the protein or that binding is dimerization dependent.

Besides the Sed5/Bos1/Bet1/Sec22 combination ( ), across cisternal maturation Sed5 forms SNARE bundles in Golgi compartments located downstream of Uso1 domains (Pelham, 1999). In fungi, membrane fusion in the medial Golgi involves the Sed5 partners Gos1 (Qb) and Sft1 (Qc) substituting for Bos1 and Bet1, respectively, but neither Gos1 nor Sft1 bound wild-type or E6K/G540 Uso1 GHD (Figure 11F), demonstrating that interaction of Uso1 with Bos1 and Bet1 is highly specific. Therefore, it seems fair to conclude that increased binding for a SNARE receptor underlies the mechanism by which mutant Uso1 bypasses the need for RAB1 in the ER/Golgi interface.

In summary, Uso1 is an essential protein acting in the ER/Golgi interface, and we report here several important findings. We show that (i) the Golgi GTPase RAB1, which is essential for viability, becomes dispensable if there is an alternative method to recruit Uso1 to Golgi membranes; (ii) the coiled-coil region of Uso1 is dispensable to sustain viability, implying that the tethering role of the protein is not essential either, consistent with the redundant roles of other Golgin tethers; (iii) the Uso1 GHD is essential for viability; (iv) the Uso1 GHD monomer, if present at suitably high levels, is sufficient to maintain viability; (v) that the Uso1 GHD is a direct and specific binder of the Qb SNARE Bos1, and a strong binder of the Qc Bet1; (vi) the mutation bypassing the need for RAB1 markedly increases the affinity of the GHD for Bos1, indicating that *rab1Δ* viability rescue by E6K/G540S Uso1 occurs because SNARE anchoring provides an alternative mode of recruitment to Golgi membranes to that provided physiologically by RAB1.

### Mechanistic insights guided by AlphaFold2 predictions

To gain further insight into the mechanisms by which the double E6K/G540S bypasses RAB1 we exploited AlphaFold2 to model the Uso1^GHD^ domain alone, or together with each of the individual SNAREs, with both Bos1 and Bet1 simultaneously, and with RAB1.

Consistent with biochemical data, in the predicted model Bet1 and Bos1 interact with a medial and a C-terminal region, respectively, of Uso1^GHD^. (Figure 12; Figure 12— supplement 1 and —supplement 2). Bet1 docks against a region of the GHD that is not affected by the mutations. In contrast, Gly540 and its environs dock against the N-terminal, triple *α*-helical Habc domain of Bos1. In all likelihood Gly540Ser mediates the increased binding of mutant Uso1^GHD^ to Bos1. AlphaFold2 predicts that Uso1^GHD^ interacts with Bos1 through a surface composed by the N-terminal part of the first Bos1 Habc *α*-helix and the loop between *α*-helices 2 and 3 (Figure 12A and B). The binding surface in Uso1^GHD^ involves the second *α*-helices of the ARM10 and ARM11 repeats, (*α*-helices 26 and 29), and the loop connecting the first two *α*-helices of ARM10 (Figure 12B). In AlphaFold2 models, Uso1 Gly504 (wild-type) is located at the beginning of Uso1^GHD^ *α*-helix 29, at the heart of the interaction surface, contributing to the Uso1-Bos1 interaction by coordinating the amide group of the Uso1 backbone with the Bos1 Glu58 carboxylate to create a hydrogen bond (Figure 12C). According to the most confident prediction, Gly504Ser results in the hydroxymethyl side chain protruding into a small pocket rimmed by the side chains of Bos1 Leu59 and Ile60, such that the Uso1 Ser504 hydroxyl group hydrogen-bonds the amide group of Bos1 Glu58 (Figure 12C). Besides creating a new hydrogen bond, Gly540Ser would increase the surface of interaction slightly, from ∼825.2 Å^2^ to ∼831.1 Å^2^, implying that these alterations, together with a minor shift in the environs of Gly540 detected by models (which might facilitate additional interactions), would strengthen the binding of the Uso1^GHD^ to the Bos1 surface.

**Figure 12.**
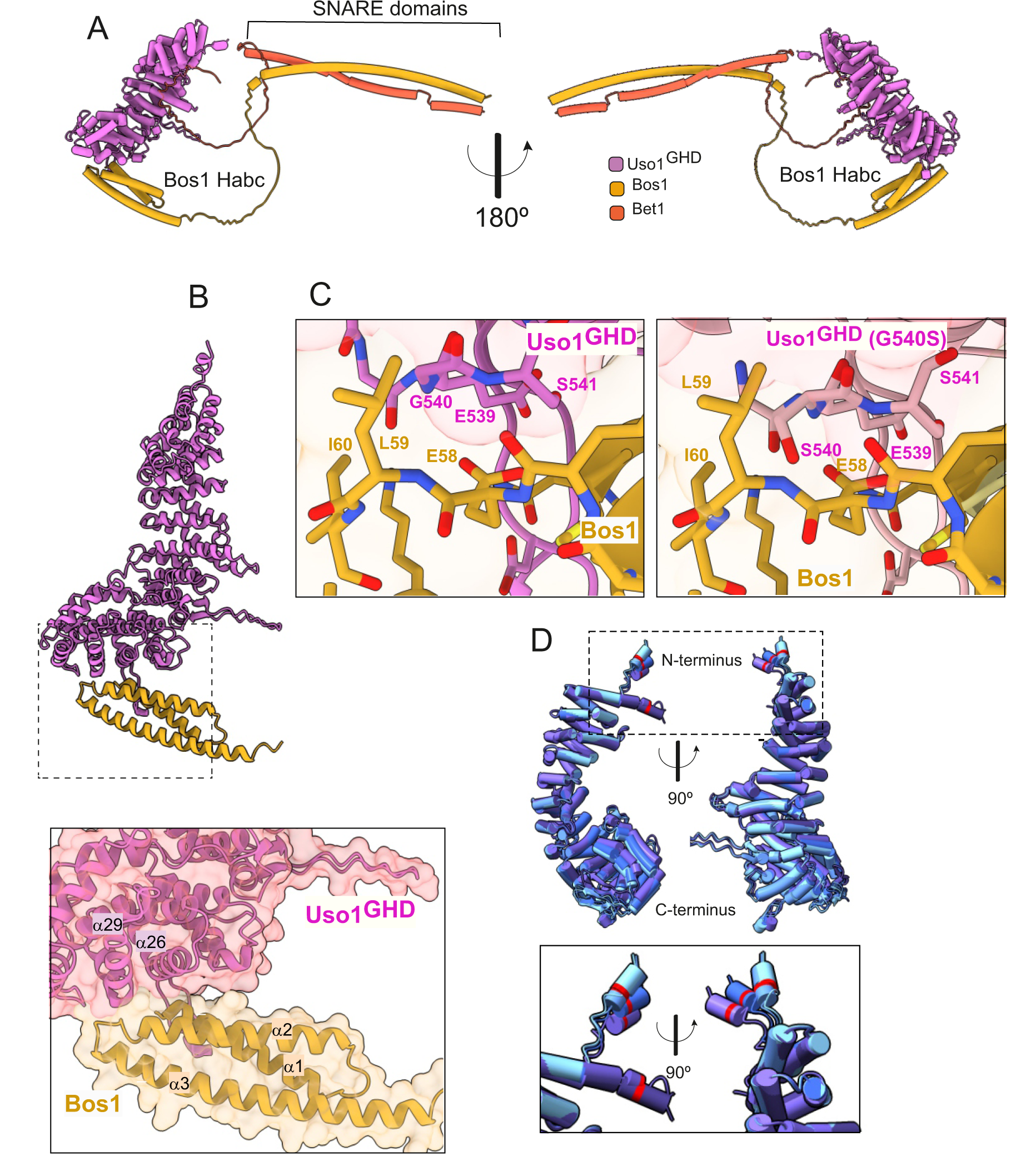
AlphaFold2 models provide insight into the additive mode of suppression shown by E6K and G540S. (A). Model of full length Uso1 bound to the ER/Golgi SNAREs Bos1 and Bet1. (B). Top, ribbon representation of the Bos1 N-terminal Habc domain and Uso1GHD. Bottom, Inset combining surface and ribbon depiction. (C). Increased binding of Bos1 to G540S Uso1 appears to involve insertion of Ser540 into a pocket located in the Habc domain of the Qb SNARE. Partial view of the Bos1- Uso1 GHD surface of interaction in the wild type (left) and mutant (right) models. G540 and S540 are annotated. (D). The N-terminal amphipathic *α*-helix of Uso1 comprising the E6K substitution lies within a flexible stretch of the protein that might facilitate its insertion into membranes. Alignment of six independent predictions, with Glu6 highlighted in red. The Uso1 GHD was modeled alone, in a complex with SNARE proteins or with Ypt1. The N-terminal a- helix (boxed) adopts different positions, suggesting high flexibility.

The mechanism by which Glu6Lys contributes to increase the recruitment of Uso1 to Golgi membranes appears to be different. This glutamate is located in a region with low pLDDT score which shows different conformations depending on the model (Figure 2D, boxed), suggesting that this region is difficult to predict due to flexibility. However, models concur in the prediction of an amphipatic N-terminal *α*-helix containing Glu6, whose substitution by Lys (as in E6K) reinforces the positive charge of this *α*-helix, which would be inserted into the vesicle membrane, potentially contributing to Uso1 recruitment to COPII vesicles

In the case of the strong Uso1^GHD^-Bet1 interaction (Figure 12—figure supplement 1), the N-terminal region of Bet1 consisting of *circa* 80 amino acids is disordered, and the pLDDT score of the different models is understandably low. However, all structural models depicting Bet1 interacting with Uso1^GHD^ (e.g. in the context of the whole SNARE complex, Bet1 alone or the isolated Bet1 N-terminal) consistently show a region where the pLDDT is higher. This region forms a kink in this N-terminal part of the Bet1 that protrudes into the surface created by the *α*-helices 13 and 16 of ARM4 and ARM5 (Figure 12—figure supplement 1). Surface representations show that this section of the Bet1 polypeptide covers ∼ 1300 Å^2^ of the GHD, docking against the same side of the boomerang-shaped solenoid as Bos1, which is consistent with the orientation that these SNAREs should take during the formation of the SNARE pin (Figure 12—figure supplement 2)

We next modelled RAB1 binding to the GHD in the absence and presence of Uso1 binders Bos1 and Bet1. AlphaFold2 predicts that the GHD interacts with RAB1 through a binding surface formed by the ARM3-5. repeats. On the other hand, the interactive region of RAB1 conforms to canons, as Uso1 *α*-helices 8 and 9 (ARM3) interact with the Switch II region while *α*-helices 11 (ARM-4) and 14 (ARM-5) interact with the Switch I) (Figure 12—figure supplement 2, A and B). Importantly, the model indicates that within the Uso1-GHD jai alai basket, RAB1 binds to the opposite (convex) side of the Bet1 interacting area at the concave side, and away from the C-terminal helix where the Bos1 Habc domain predictably binds, which would allow Uso1 to bind these three interactors simultaneously (Figure 12—figure supplement 2C). In addition, the predicted models supported two highly suggestive but as yet speculative implications. One is that the position of the RAB1 hypervariable domain, which anchors the GTPase to the membrane through is prenylated C-terminus, is compatible whith the hypothetical membrane binding of the N-terminal Uso1 amphipatic helix (Figure 12—figure supplement 2D); The second is that Uso1-RAB1 would be in an orientation that facilitates the docking of RAB1- loaded ER-derived vesicles with an acceptor membrane where SNARE zippering occurs (see discussion); this orientation implies that the donor membrane (the position of the N-terminal Uso1 helix) and the acceptor membrane (the TMDs of the SNAREs) is compatible with the Uso1 CTR contributing to tethering through interaction with Bug1/Grh1.

## Discussion

RAB1 regulates transport at the ER/Golgi interface. Using an unbiased forward genetic screen to identify subordinated genes accounting for its essential role, we isolated two extragenic mutations resulting in single-residue substitutions in the RAB1 effector Uso1, usually regarded as a tether. The single-residue substitutions lie at opposite ends of the jai alai basket-shaped GHD; both are individually able to rescue viability of *rab1Δ* strains at 30°C and, when combined, even at 42°C. Subcellular localization experiments hinted at the mechanism by which the double mutation rescues *rab1*Δ lethality. Uso1 plays its physiological role on an early Golgi compartment, where it largely colocalizes with RAB1, with relocation to the cytosol in a RAB1-deficient background. Under normal circumstances, the Uso1 CTR acts in concert with RAB1 to recruit the protein to the Golgi. Genetic evidence (Figure 8) showed that this contribution requires the Golgi-localized tether composed of the membrane anchor Grh1 and its associated golgin BUG1 (Behnia et al., 2007), homologues of human GRASP65 and GM130, respectively. In the absence of RAB1, engagement of the CTR with BUG1 is insufficient to stabilize wild-type Uso1 on membranes. However, the double E6K/G540S substitution relocalizes Uso1 to Golgi structures, suggesting that mutant Uso1 GHD had gained affinity for another element, thereby compensating for the loss of RAB1.

By S-tag co-precipitation experiments we identified proteins associating with wild-type and E6K/G540S Uso1 baits. Prominent among these were the four SNAREs, Sed5, Bos1, Bet1 and Sec22, mediating fusion events at the ER/ Golgi interface, and the SNARE regulator Sec18, indicating that Uso1 is a component of the SNARE fusion machinery. This was suggested by previous studies with *S. cerevisiae* showing that overexpression of Bet1, Bos1 and Sec22 suppresses *uso1-1* and that of Sec22 and Bet1 weakly suppresses *uso1Δ* (Sapperstein et al., 1996). Moreover, in mammalian cells crosslinking studies with p115 identified Sed5, membrin (Bos1) and mBet1 as its weak interactors (Allan et al., 2000), suggesting that contributing to the ER/Golgi SNARE machinery is a conserved feature of Uso1/p115 family members. Of the above four *A. nidulans* SNAREs, Bet1 and Bos1, which co-precipitated weakly with the wild-type, were dramatically enriched with the mutant. Pull-down assays showed that full-length *Aspergillus* Uso1 interacts weakly with Sec22 and strongly with Bet1, irrespective of whether the bait was wild-type or E6K/G540S, indicating that they are direct physiological interactors. In sharp contrast, the interaction of E6K/G540S Uso1 with the Qb Bos1 was markedly augmented, strongly suggesting that this increase contributes substantially to bypass RAB1. Both the high levels of binding to Bet1, and the marked increase in binding to Bos1 resulting from E6K/G540S were tracked down to the GHD, which is sufficient to bind these SNAREs.

AlphaFold2 models predicting the regions of interaction of the GHD with Bos1 and Bet1 showed that, of the two Uso1 residue substitutions, only Gly540Ser maps to the region mediating the interaction with Bos1, whereas neither affected the interacting region with Bet1, agreeing with GST pull-downs. Interaction with this SNARE is predicted to involve a large surface, consistent with the strong “constitutive” binding of Bet1 with the GHD. AlphaFold2 also predicted that RAB1 binds to the convex face of the GHD solenoid, in a position that would permit the simultaneous binding of Qb and Qc SNARES and the GTPase. In addition, AlphaFold2 also detected a previously unnoticed amphipathic *α*- helix in the N-terminal region of Uso1. The Glu6Lys substitution falls within this helix, making its global positive charge even greater, strongly suggesting that this substitution increases Uso1 GHD binding to membranes. That the two residue substitutions act by different mechanisms is coherent with their showing additivity to suppress RAB1 deficit.

The Uso1/p115 family has been implicated in the regulation of SNARE complexes (Allan et al., 2000). (Shorter et al., 2002) first reported that p115 binds to SNAREs and proposed that they would ‘catalyze’ the formation of the ER/Golgi SNARE bundle. The direct interaction of p115 with unassembled mBet1 and Sec22 agrees with this role (Wang et al., 2014). However, there are significant gaps in this model: Uso1 overexpression suppresses a partial deficit of *ypt1/RAB1*, but not *ypt1*Δ*, whereas* Ypt1 overexpression suppresses *uso1Δ*, indicating that Uso1 acts and in concert or upstream of RAB1. [Of note, overexpression of Uso1 also suppresses *bet3-1*, a *ts* allele inactivating the TRAPPIII complex, which is the RAB1 GEF (Galindo et al., 2021; Jiang et al., 1998; Pinar et al., 2019; Riedel et al., 2017; Thomas et al., 2018)]. The grid of reported genetic interactions strongly indicates that RAB1 regulates SNAREs (Brandon et al., 2006; Lupashin and Waters, 1997; Sapperstein et al., 1996). Notably, our viability rescue experiments show that E6K/G540S bypasses this non-canonical, yet essential role of RAB1. As the double mutant substitution increases the recruitment of USO1 to the SNARE machinery and as E6K/G540S suppresses *rab1Δ*, the most parsimonic interpretation of the data is that the role of RAB1is cooperating to recruit Uso1 to the SNARE complexes, a cooperation that is no longer needed when recruitment is ensured by other means. That p115 regulates SNAREs’ assembly was proposed by (Shorter et al., 2002). However, a fundamental difference with our conclusions is that they attributed this role to the CCD, whereas we identify here the GHD as the positively-acting player.

We also note that our experiments provide a mechanistic interpretation as to why overexpression of Bet1, Bos1, Sec22 and Ypt1/RAB1 rescue the viability of *uso1-1* removing a substantial portion of the yeast Uso1 CCD (Sapperstein et al., 1996; Seog et al., 1994). A similar *Aspergillus* allele results in marked protein instability (Figure 9). Thus, in all likelihood, suppression by overexpression of known direct interactors involves stabilization of the *uso1-1* product.

In summary, our work firmly establishes that the essential role of Uso1 resides not in its CCD domain tethering donor and acceptor membranes, but in the GHD. When expressed at sufficient levels this domain is capable of fully complementing *uso1Δ*, which is definitive evidence that neither the tethering function of Uso1 nor dimerization (the GHD is monomeric) is required for the protein to play its essential role. As the GHD binds two of the four SNAREs of the bundle, the simplest interpretation is that the essential role of Uso1 is regulating SNAREs.

### Ideas and Speculation

While the molecular details of this regulation will be addressed in future, it is tempting to speculate that the GHD contributes, with Sly1, to orientate SNAREs to form a productive bundle, acting as chaperones, similarly to the HOPS SM component Vps33, which appears to align SNAREs in a pre-zippering stage, facilitating their assembly (Baker and Hughson, 2016; Baker et al., 2015; Ren et al., 2009; Yu and Hughson, 2010; Zhang and Hughson, 2021; Zhang and Yang, 2020). A second speculative interpretation is that tethering occurs in two steps, with golgins acting at long distances, approximating vesicles to the vicinity of SNAREs. Then SNAREs would engage RAB1–Uso1 to serve as short-range tether preceding the zippering up of membranes. The > 700 Å-long Uso1 CCD would cooperate with Bug1 in the first step, whereas the Uso1 GHD would cooperate with RAB1 and Bet1/Bos1 in the second, exploiting the fact that GHD binders use surfaces located at opposite sides of the *α*-solenoid (Figure 12—figure supplement 2). Such arrangement implies that RAB1 C-terminal isoprenoids inserting into the donor vesicle membrane would be *circa* 220 Å apart from the C-termini of the SNAREs inserted in the acceptor membrane. Two-step tethering might impose one additional level of specificity, preventing unproductive fusion events mediated by other SNARES circulating through the ER/ interface and directing, by way of Uso1 interactions, incoming vesicles to fusion-competent areas enriched in target SNAREs (Bentley et al., 2006).

## Materials and Methods

### Aspergillus techniques

Standard *A. nidulans* media were used for growth tests, strain maintenance and conidiospore harvesting (Cove, 1966). GFP-, HA3- and S-tagged alleles were introduced by homologous recombination-mediated gene replacement, using transformation (Tilburn et al., 1983) of recipient *nkuAΔ* strains deficient in the non-homologous end joining pathway (Nayak et al., 2005). Complete strain genotypes are listed in supplemental table I. These alleles were usually mobilized into the different genetic backgrounds by meiotic recombination (Todd et al., 2007).

Null mutant strains were constructed by transformation-mediated gene replacement, using as donor DNA cassettes made by fusion PCR (primers detailed in supplemental table II) carrying appropriate selectable markers (Szewczyk et al., 2006). Integration events were confirmed by PCR with external primers. When allele combinations were expected to be synthetically lethal or severely debilitating, the corresponding strains were constructed by sequential transformation, but the second such manipulation was always carried out using *pyrG^Af^* as selective marker, which favors the formation of heterokaryons in which untransformed nuclei supported growth (Osmani et al., 1988). Conidiospores, in which single nuclei had segregated (i.e., homokaryotic nuclei), were scrapped and streaked onto plates carrying doubly-selective medium. Absence of growth or appearance of microcolonies, combined with a positive PCR diagnostic of heterokaryosis of the primary transformants, was taken as indication of lethality. Whenever possible, colony PCR of microcolonies was always used to genotype the desired genetic intervention

The following proteins were C- or N-terminally tagged endogenously, using cassettes constructed by fusion PCR (Nayak et al., 2005; Szewczyk et al., 2006): Uso1-GFP and Uso1^E6K/G540S^-GFP, Uso1-HA3 and Uso1^E6K/G540S^-HA3; Uso1-S and Uso1^E6K/G540S^-S; BapH-S (Pinar and Peñalva, 2017), Sec13-mCherry (Bravo-Plaza et al., 2019; Hernández-González et al., 2019), Gea1-mCherry and Sec7-mCherry (Arst et al., 2014), mCherry-Sed5 (Pantazopoulou and Peñalva, 2011), mCherry-RAB1 (Pinar et al., 2013), HA3-Sed5, HA3-Bet1, HA3-Bos1, HA3-Sec22, Sec18-HA3, Grh1-HA3, Bug1-HA3, Coy1-HA3, COG2-HA3, and *β*-COP-HA3.

### Antibodies for western blotting

| Antibody | dilution | Origin | Reference |
| --- | --- | --- | --- |
| Primary |  |  |  |
| α-HA | 1:1000 | rat | 3F10 clone, Roche |
| α-His6x tag | 1:10 000 | mouse | #631212, Clontech |
| α-tubulin | 1:5000 | mouse | DM1A clone, Sigma |
| α-Uso1 | 1:1000 | rabbit | Polyclonal antiserum, Davids Biotechnologie |
| Secondary (HRP-conjugated) |  |  |  |
| α-rat IgG | 1:4000 | goat | #3010-05, Southern Biotech |
| α-mouse IgG | 1:5000 | goat | #A9044, Sigma |
| α-rabbit IgG | 1:2000 | donkey | #NA934, Amersham |

Antiserum against Uso1 was raised in rabbits by Davids Biotechnology. Animals were immunized with the Uso1 GHD (residues 1-659), tagged with His6x. Recombinant expression in *E. coli* and Ni^2+^ affinity purification is described below. Target antibodies were purified from raw antiserum by affinity chromatography through Hi-Trap NHS columns (#17-0716-01, Cytiva) charged with Uso1 antigen following the manufacturer’s instructions. Affinity-bound antibodies were eluted with 100 mM glycine (pH 3.0), then neutralized with 2 M Tris to a pH of 7.5 and stored at -20°C.

### *inuA* promoter-driven expression of Uso1 GHD in *Aspergillus*

The Uso1 GHD was expressed in a sucrose-inducible manner from an *in locus* replacement of the *inuA* gene (AN11778) ORF encoding inulinase by the GHD coding sequence, such that its expression was driven by the *inuA* promoter, which is induced on sucrose and non-induced on glucose (Hernández-González et al., 2018). The gene replacement cassette was assembled through fusion PCR of 4 different elements, listed here from 5’ to 3’: (1) inuA promoter, (2) cDNA sequence encoding Uso1 wt or mutant E6K/G540S GHD (residues 1 – 659), (3) *Aspergillus fumigatus riboB* gene as selection marker and (4) *inuA* gene 3’-flanking region. A *pyrG*89, *nkuA*Δ::bar, *riboB*2 *A. nidulans* strain was transformed with this cassette, replacing the inuA gene. The resulting strain was subsequently transformed with a *uso1*Δ deletion cassette (*A. fumigatus pyrG* as selection marker) to ablate endogenous of Uso1, such that the GHD was the only moiety of Uso1 present.

### Plasmids for protein expression in *E. coli*

#### (I) His6x-tagging constructs

<u>pET21b-Uso1-His6x</u> and <u>pET21b-Uso1(E6K/G540S)-His6x</u>: cDNA encoding full length Uso1 (residues 1 -1103) was cloned into a pET21b *Nde*I/*Not*I linearized vector.

<u>pET21b-Uso1ΔCTR-His6x</u> and <u>pET21b-Uso1(E6K/G540S)ΔCTR-His6x</u>: lacking the C- Terminal Region of the Coiled Coil Domain (residues 1 – 1040)

<u>pET21b-Uso1GHD-His6x</u> and <u>pET21b-Uso1(E6K/G540S)GHD-His6x</u>: cDNA encoding the Globular Head Domain of Uso1 (residues 1 - 659) was cloned as a *Nde*I/*Xho*I insert into a pET21b *Nde*I/*Xho*I linearized vector.

<u>pET21b-Uso1 CCD-His6x</u>: cDNA encoding Uso1 Coiled Coil Domain (residues 660- 1103) was cloned as a *Nde*I/*Not*I insert into a pET21b *Nde*I/*Not*I linearized vector.

### TNT^®^ expression plasmids

<u>pSP64-Sly1-HA</u>: this plasmid carries cDNA encoding full length Sly1 (AN2518) C- terminally tagged with a HA3x epitope, cloned as an *Nsi*I/*Sac*I insert into *Pst*I/*Sac*I pSP64(PolyA) vector

### GST-tagging constructs

<u>pET21b-Sed5-GST:</u> cDNA encoding Sed5/AN9526 cytoplasmic domain (residues 1–322) C-terminally tagged with GST, was cloned as a *Nde*I/*Sal*I insert into a pET21b *Nde*I/*Xho*I linearized vector.

<u>pET21b-Bos1-GST:</u> cDNA encoding Bos1/AN11900 cytoplasmic domain (residues 1–219) C-terminally tagged with GST, was cloned as a *Nde*I/*Sal*I insert into a pET21b *Nde*I/*Xho*I linearized vector.

<u>pET21b-Bet1-GST:</u> cDNA encoding Bos1/AN5127 cytoplasmic domain (residues 1–71) C-terminally tagged with GST, was cloned as a *Nde*I/*Sal*I insert into a pET21b *Nde*I/*Xho*I linearized vector.

<u>pET21b-Sec22-GST:</u> cDNA encoding Sec22/ ASPND00903 cytoplasmic domain (residues 1–198) C-terminally tagged with GST, was cloned as a *Nhe*I/*Sal*I insert into a pET21b *Nhe*I/*Xho*I linearized vector.

<u>pET21b-Sso1-GST:</u> cDNA encoding Sso1/AN3416 cytoplasmic domain (residues 1–271) C-terminally tagged with GST, was cloned as a *Nde*I/*Sac*I insert into a pET21b *Nde*I/*Sac*I linearized vector.

<u>pET21b-Gos1-GST:</u> cDNA encoding Gos1/AN1229 cytoplasmic domain (residues 1–208) C-terminally tagged with GST, was cloned as a *Nde*I/*Sal*I insert into a pET21b *Nde*I/*Xho*I linearized vector.

<u>pET21b-Sft1-GST:</u> cDNA encoding Sft1/AN10508 cytoplasmic domain (residues 1–73) C-terminally tagged with GST, was cloned as a *Nde*I/*Sal*I insert into a pET21b *Nde*I/*Xho*I linearized vector.

### Co-precipitation experiments with total cell extracts

Preparation of *Aspergillus* total cell extracts was done as described, with minor modifications (Pinar et al., 2019) (Pinar et al., 2019). 70 mg of lyophilized mycelium were ground with a ceramic bead in a Fast Prep (settings: 20 sec, power 4). The resulting fine powder was resuspended in 1.5 ml of extraction buffer [25 mM HEPES-KOH (pH 7.5), 200 mM KCl, 4 mM EDTA, 1% (v/v) IGEPAL CA-630 (NP-40 substitute, #I8896, Sigma), 1 mM DTT, 2 μM MG-132 proteasome inhibitor (#S2619, SelleckChem) and cOmplete**^®^** ULTRA EDTA-free inhibitor cocktail (#5892953001, Roche). Approximately 0.1 ml of 0.6 mm glass beads were added and thoroughly mixed. This suspension was homogenized with a 10 sec pulse at the Fast Prep (power 6) followed by a 10 min incubation at 4°C. This homogenization step was repeated two times before clarifying the extract by centrifugation at 4°C and 15,000 × *g* in a microcentrifuge. Total protein concentration of the extracts was determined by Bradford. Bovine Serum-Albumin BSA was then added as a blocking agent to the cell extract (final concentration 1% (w/v)). Binding reactions were carried out in 0.8 ml Pierce centrifuge columns (#89869, ThermoFisher): 9 mg of protein were mixed with 20 µL of S-protein Agarose beads (#69704, Novagen), that had been previously washed in extraction buffer with 1% (w/v) BSA. This buffer was also added to complete final reaction volume of 0.6 ml. The mix was incubated for 3 h at 4°C in a rotating wheel. Columns were then opened at the bottom and gently centrifuged to remove the supernatant and collect the protein-bound beads. These were resuspended in extraction buffer without inhibitors and incubated in rotation for 10 min at 4°C, followed by two more washing steps in extraction buffer without detergent and inhibitors. To elute proteins bound to the beads, 30 µl of Laemmli loading buffer [62.5 mM Tris-HCl (pH 6.0), 6 M urea, 2% (w/v) SDS and 5% (v/v) β-mercaptoethanol] were added and the columns incubated at 90°C for 2 min. The columns were centrifuged to collect the eluate, of which a 40% of the final volume were resolved in a SDS-polyacrylamide gel and then transferred to nitrocellulose for α-HA (#3F10, Roche) western blotting.

### Purification of Uso1 constructs tagged with His6x

Full-length His6-tagged Uso1, Uso1ΔCTR, Uso1 GTD and Uso1 CCD constructs, wild type and mutant versions, were expressed in *E. coli* BL21(DE3) cells harboring pET21b- His6 derivatives and pRIL. Bacteria were cultured at 37°C in LB medium containing ampicillin and chloramphenicol until reaching an OD_600nm_ of 0.6. Then, IPTG was added to a final concentration of 0.1 mM. Cultures were shifted to 15°C and incubated for 20 h.

Bacterial cells were collected by centrifugation and pellets stored at -80°C. For purification, frozen pellets were thawed in ice and resuspended in ice-cold bacterial cell lysis buffer [20 mM sodium phosphate buffer, pH 7.4, 500 mM KCl, 30 mM imidazole, 5% (v/v) glycerol, 1 mM β-mercaptoethanol, 1 mM MgCl_2_, 0.2 mg/ml lysozyme and 1 μg/ml of DNAse I and cOmplete^®^ protease inhibitor cocktail (#11873580001, Sigma)]. This cell suspension was mechanically lysed in a French press (1500 kg/cm^2^) and the resulting lysate was centrifuged at 10,000 × *g* and 4°C for 20 min to remove the cell debris. The supernatant was then transferred to polycarbonate tubes and centrifuged at 100,000 × *g* and 4°C for 1 h in a XL-90 ultracentrifuge (Beckman Coulter) . 50 ml of cleared lysate were incubated with 400 μL of Ni-Sepharose High Performance beads (#17526801, Cytiva) for 2 h at 4°C. After this step, His-tagged protein-bound beads were pelleted at low-speed centrifugation and washed three times in lysis buffer [20 mM sodium phosphate buffer, pH 7.4, 500 mM KCl, 5% (v/v) glycerol, 1 mM β- mercaptoethanol] with increasing concentrations of imidazole. Finally, Ni2+bound His6 proteins were eluted 0.5 M imidazole buffer. 5 ml of eluted protein were loaded onto a HiLoad 16/600 Superdex 200 column (Cytiva) and run at 1 ml/min flow rate on an AKTA HPLC system, using phosphate buffered saline PBS containing 5% (v/v) glycerol and 1 mM β-mercaptoethanol. Fractions containing protein were pooled, analyzed for purity by SDS-PAGE followed by Coomassie staining, and finally quantified on a UV-Vis spectrophotometer before being stored at -80°C.

### Purification of SNARE constructs tagged with GST

cDNAs encoding the cytosolic domains of SNAREs fused to a C-terminal GST were cloned into pET21b. Bacterial cultures and protein expression conditions were as described above for Uso1-His6 constructs. Frozen pellets were thawed in ice and resuspended in chilled bacterial cell lysis buffer [25 mM Tris-HCl (pH 7.4), 300 mM KCl, 5 mM MgCl_2_, 1 mM DTT, 0.5 mg/ml lysozyme, 1 μg/ml of DNAse I and cOmplete^®^ protease inhibitor cocktail (#11873580001, Sigma)]. This cell suspension was incubated for 30 min in ice before being mechanically lysed in a French press (1500 kg/cm^2^). The lysate was incubated for a further 30 min on ice and centrifuged at 20,000 × *g* and 4°C for 30 min. After adding 10 mM EDTA to the clarified supernatant to stop DNAse I activity, it was transferred to a 50 ml tube, mixed with 500 µL of glutathione Sepharose beads 4B (#17075601, Cytiva) and rotated for 2 h at 4°C. After incubation, SNARE-GST-bound beads were pelleted by gentle centrifugation and washed three times for 10 min at 4°C in 25 mM Tris-HCl (pH 7.4), 500 mM KCl, 5 mM EDTA, 1 mM DTT and subsequently transferred to a 0.8 ml Pierce column. Beads were washed 6 times (10 min at RT) in 200 µL of elution buffer [50 mM Tris-HCl (pH 8.0), 200 mM KCl, 10 mM glutathione and 1 mM DTT]. These fractions were collected and pooled (∼1 ml), then buffer-exchanged to storage buffer (PBS, 5% (v/v) glycerol and 0.1 mM DTT) in a PD MidiTrap G-25 column. Protein concentration and purity was assessed by spectrophotometry and SDS-PAGE followed by Coomassie staining. Protein stocks were kept frozen at -80°C.

### SNARE-GST pull-downs with purified Uso1-His6 constructs (Uso1, Uso1 GHD)

Binding reactions were performed in 0.8 ml Pierce centrifuge columns. 75 µg of purified SNARE-GST were mixed with 15 µL of glutathione Sepharose 4B beads and storage buffer to a final volume of 0.3 ml. Columns were rotated at 4°C for 2 h before the supernatant was removed after low speed centrifugation. Subsequently, His6 preys were added to a final concentration of 0.2 µM in a total volume of 0.4 ml of pull-down binding buffer [25 mM HEPES-KOH (pH 7.5), 150 mM NaCl, 10% (v/v) glycerol, 0.1% (v/v) Triton X-100 and 0.1 mM DTT]. Columns were rotated overnight at 4°C. Beads were collected by gentle centrifugation and washed three times for 10 min with ice-cold binding buffer, before eluting bound proteins with 30 µL of Laemmli loading buffer pre-heated at 90°C. 0.5 % of the samples (eluted material or flow-through) were run onto 8% SDS-polyacrylamide gels that were transferred to nitrocellulose membranes which were reacted with α-His tag antibody (#631212, Clontech). Quantitation of band intensities was done with ImageLab software (BioRad). In parallel, 4% of the elution sample volume was loaded onto 10% SDS-polyacrylamide gel and stained with coomassie dye (BlueSafe, NZY) to confirm recovery of SNARE-GST baits.

### Pull-down of TNT^®^-expressed Sly1-HA3

Sly1-HA3 was synthesized with the TNT^®^ SP6 Quick Coupled Transcription/Translation system (#L2080, Promega), according to the instructions of the manufacturer. The reaction was primed with 1 μg of pSP64::Sly1-HA3 cDNA. 10 μL of the resulting mix were combined with 15 µL of glutathione-Sepharose beads, previously loaded with SNARE-GST baits as described above, in 0.4 ml of pull-down binding buffer, using 0.8 ml Pierce columns that were rotated overnight at 4°C before beads and flow-through were recovered after gentle centrifugation. Beads were washed three times for 10 min at 4°C in pull-down binding buffer before eluting bound material with 30 µL of Laemmli loading buffer for 2 min at 90°C. 20% of the elution sample volume was analyzed by western blotting with α-HA tag antibody (#3F10, Roche) for Sly1-HA immunodetection.

### Size exclusion chromatography of HA-tagged cell extracts

Gel filtration experiments were performed as described (Bravo-Plaza et al., 2019). Briefly, 200 µL of cell extract were loaded onto a Superose 6 10/300 column (Pharmacia) equilibrated with running buffer [25 mM Tris-HCl (pH 7.5), 600 mM KCl, 4 mM EDTA, 1 mM DTT]. Fractions of 0.5 ml were collected, from which 80 µL were mixed with 40 µL of Laemmli loading buffer and denatured at 90°C. 25 µL of these samples were resolved by SDS-PAGE and analyzed by western blotting with α-HA3 tag antibody (#3F10, Roche) for western blotting. Sizing standards were myoglobin (17 kDa), BSA (67 kDa), aldolase (158 kDa) ferritin (449 kDa), thyroglobulin (669 kDa) and dextran blue (Vo).

### Analytical ultracentrifugation: sedimentation velocity

Sedimentation velocity assays and subsequent raw data analysis were performed in the Molecular Interactions Facility of the Centro de Investigaciones Biológicas Margarita Salas. Samples (320 µL) in PBS containing 5% (v/v) glycerol and 1 mM β- mercaptoethanol were loaded into analytical ultracentrifugation cells, which were run at 20°C and 48,000 rpm in a XL-I analytical ultracentrifuge (Beckman-Coulter Inc.) equipped with UV-VIS absorbance and Raleigh interference detection systems, using an An-50Ti rotor, and 12 mm Epon-charcoal standard double-sector centerpieces. Sedimentation profiles were recorded at 230 nm. Differential sedimentation coefficient distributions were calculated by least-squares boundary modelling of sedimentation velocity data using the continuous distribution *c(s)* Lamm equation model as implemented by SEDFIT(Schuck, 2000). Experimental Svedberg coefficient values were corrected to standard conditions (*s_20,w_* : water, 20°C, and infinite dilution) using SEDNTERP software

### Dynamic Light Scattering, DLS

DLS experiments were carried out in a Protein Solutions DynaPro MS/X instrument at 20C using a 90° light scattering cuvette. DLS autocorrelation functions, average of at least 18 replicates, were collected with Dynamics V6 software. Analysis evidenced in most cases the presence of two diffusing species, one with faster diffusion corresponding to a discrete major species and a second with substantially slower diffusion, corresponding to higher order species, with a minor contribution to the whole population within the sample. Exceptions were the globular domain GHD that appeared as a single species, and the coiled-coil construct with a larger contribution of the higher order species. Dynamics software was also employed to export the data as text files for parallel analysis using user-written scripts and functions in MATLAB (Version 7.10, MathWorks, Natick, MA). A double exponential decay model was fit to the data *via* nonlinear least squares, using as starting values the translational diffusion coefficients and relative amounts of the two different species, using as starting values those from the regularization analysis and their masses (those of the discrete species as estimated by the Svedberg equation). These values were compatible with the experimental data, rendering a similar best-fit value of the diffusion coefficient for the major species and allowing to assess its probability distribution.

### Estimate of molar mass from hydrodynamic measurements

The apparent molar masses of Uso1 and its mutants were calculated via the Svedberg equation, using the *s*- and *D*-values of the major species independently measured by sedimentation velocity and DLS, respectively.

### Fluorescence Microscopy

*A. nidulans* hyphae were cultured in Watch Minimal Medium WMM (Peñalva, 2005). Image acquisition equipment, microscopy culture chambers and software have been detailed (Pinar et al., 2022; Pinar and Peñalva, 2020). Simultaneous visualization of green and red emission channels was achieved with a Gemini Hamamatsu beam splitter coupled to a Leica DMi8 inverted microscope. Z-Stacks were deconvolved using Huygens Professional software (version 20.04.0p5 64 bits, SVI). Images were contrasted with Metamorph (Molecular Devices). Statistical analysis was performed with GraphPad Prism 8.02 (GraphPad). Uso1-GFP time of residence in cisternae was estimated from 3D movies consisting of middle planes with 400 photograms at 2 fps time resolution. Each Uso1 puncta considered in the analysis was tracked manually with 3D (x,y,t) representations generated with Imaris software (Oxford Instruments) combined with direct observation of photograms in movies and kymograph representations traced across >25 px-wide linear ROI covering the full width of the hyphae.

### AlphaFold predictions

AlphaFold2 (Jumper et al., 2021) predictions were run using versions of the program installed locally and on ColabFold (Mirdita et al., 2022) with the AlphaFold2_advanced.ipynb notebook and the MMseqs2 MSA option. In all cases, the five solutions predicted by AlphaFold2 by default were internally congruent, and we always chose the one ranked first by the software. Uso1 GHD (1-674), Bos1, Bet1 and RAB1 were initially submitted as hetero-oligomers. Subsequently, predictions were submitted as 1:1 complexes of Uso1 GHD with Bos1, Bet1 and RAB1, as described in the table below. The solutions were also very similar when comparing the different combinations displayed in the table were fed to the software, strongly supporting the validity of the results.

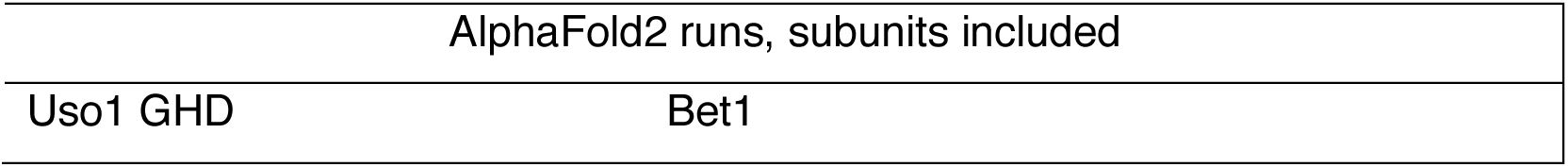
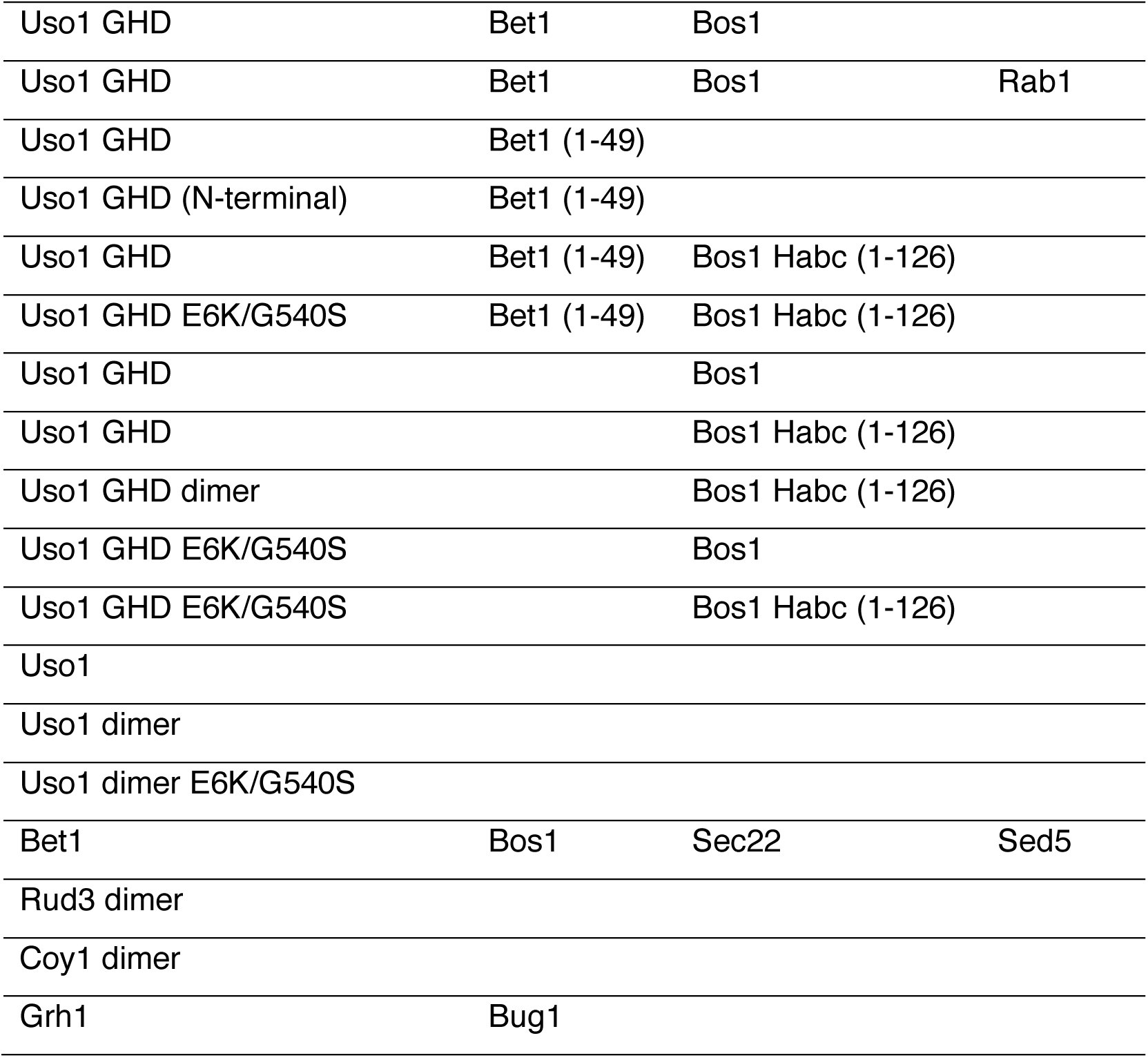

| AlphaFold2 runs, subunits included |  |
| --- | --- |
| Uso1 GHD | Bet1 |

|  |  |  |  |
| --- | --- | --- | --- |
| Uso1 GHD | Bet1 | Bos1 |  |
| Uso1 GHD | Bet1 | Bos1 | Rab1 |
| Uso1 GHD | Bet1 (1-49) |  |  |
| Uso1 GHD (N-terminal) | Bet1 (1-49) |  |  |
| Uso1 GHD | Bet1 (1-49) | Bos1 Habc (1-126) |  |
| Uso1 GHD E6K/G540S | Bet1 (1-49) | Bos1 Habc (1-126) |  |
| Uso1 GHD |  | Bos1 |  |
| Uso1 GHD |  | Bos1 Habc (1-126) |  |
| Uso1 GHD dimer |  | Bos1 Habc (1-126) |  |
| Uso1 GHD E6K/G540S |  | Bos1 |  |
| Uso1 GHD E6K/G540S |  | Bos1 Habc (1-126) |  |
| Uso1 |  |  |  |
| Uso1 dimer |  |  |  |
| Uso1 dimer E6K/G540S |  |  |  |
| Bet1 | Bos1 | Sec22 | Sed5 |
| Rud3 dimer |  |  |  |
| Coy1 dimer |  |  |  |
| Grh1 | Bug1 |  |  |

## Acknowledgements

We thank Juan R. Luque (Molecular Interactions Facility, Centro de Investigaciones Biológicas) for his help with the analytical ultracentrifuge experiments, Manuel Sánchez- Berges for HA3-tagged SNARE strains, Sara Abib and Elena Reoyo for skillful technical assistance. Thanks are due to Spain’s Ministerio de Ciencia e Innovación for grants RTI2018-093344-B100 (MAP) and predoctoral contract BES-2016-077440 (IB.-P/MAP), and to the Comunidad de Madrid for grant S2017/BMD-3691 (MAP). Grants were co- funded by European Regional Development and European Social Funds. The authors declare that they do not have any competing financial interests.

## Additional information

### Availability statement

All DNA molecules used here may reconstructed by standard techniques using primers listed in supplemental Table II. All strains listed under supplemental Table I are available for academic purposes upon reasonable request to the corresponding author. They are deposited and maintained by the corresponding laboratory in the so-denoted Madrid (MAD) collection.

## Funding

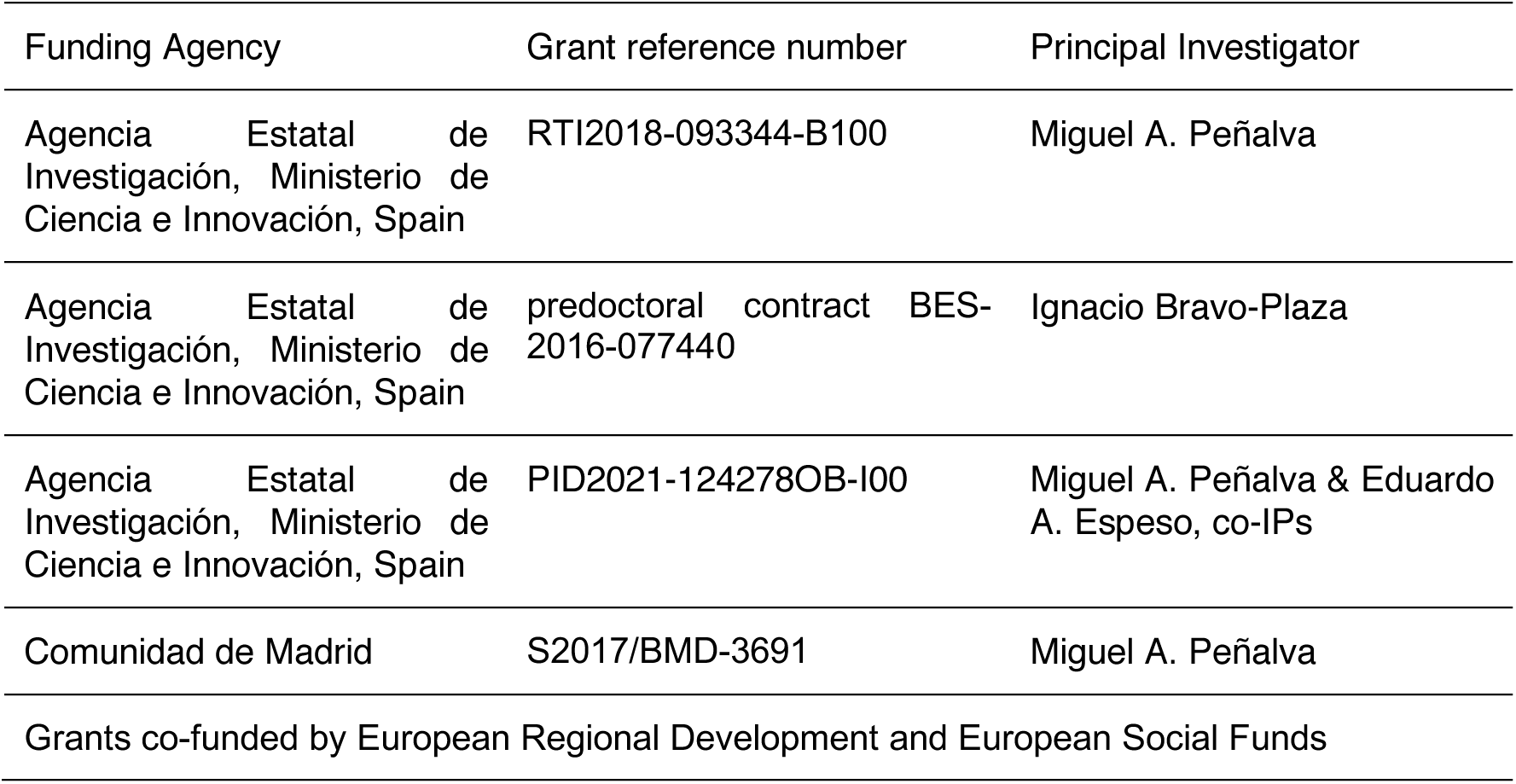

### Author contributions

Ignacio Bravo Plaza, Conceptualization, Data curation, Formal analysis, Validation, Investigation, Visualization, Writing — review and editing; Víctor G. Tagua, Data curation, Formal analysis, Validation, Investigation; Herbert N. Arst, Jr., Conceptualization, Data curation, Formal analysis, Validation, Investigation, Supervision, Writing — review and editing; Ana M. Alonso, Data curation, Formal analysis, Validation, Investigation; Mario Pinar, Conceptualization, Data curation, Formal analysis, Validation, Investigation, Supervision, Writing — review and editing ; Begoña Monterroso, Conceptualization, Data curation, Formal analysis, Validation, Supervision, Writing — review and editing; Antonio Galindo, Conceptualization, Data curation, Formal analysis, Validation, Investigation, Visualization, Writing — review and editing and Miguel Á. Peñalva, Conceptualization, Data curation, Formal analysis, Validation, Investigation, Supervision, Funding acquisition, Visualization, Writing – original draft, Writing — review and editing.

## Legends to figure

**Figure 1—figure supplement 1:**
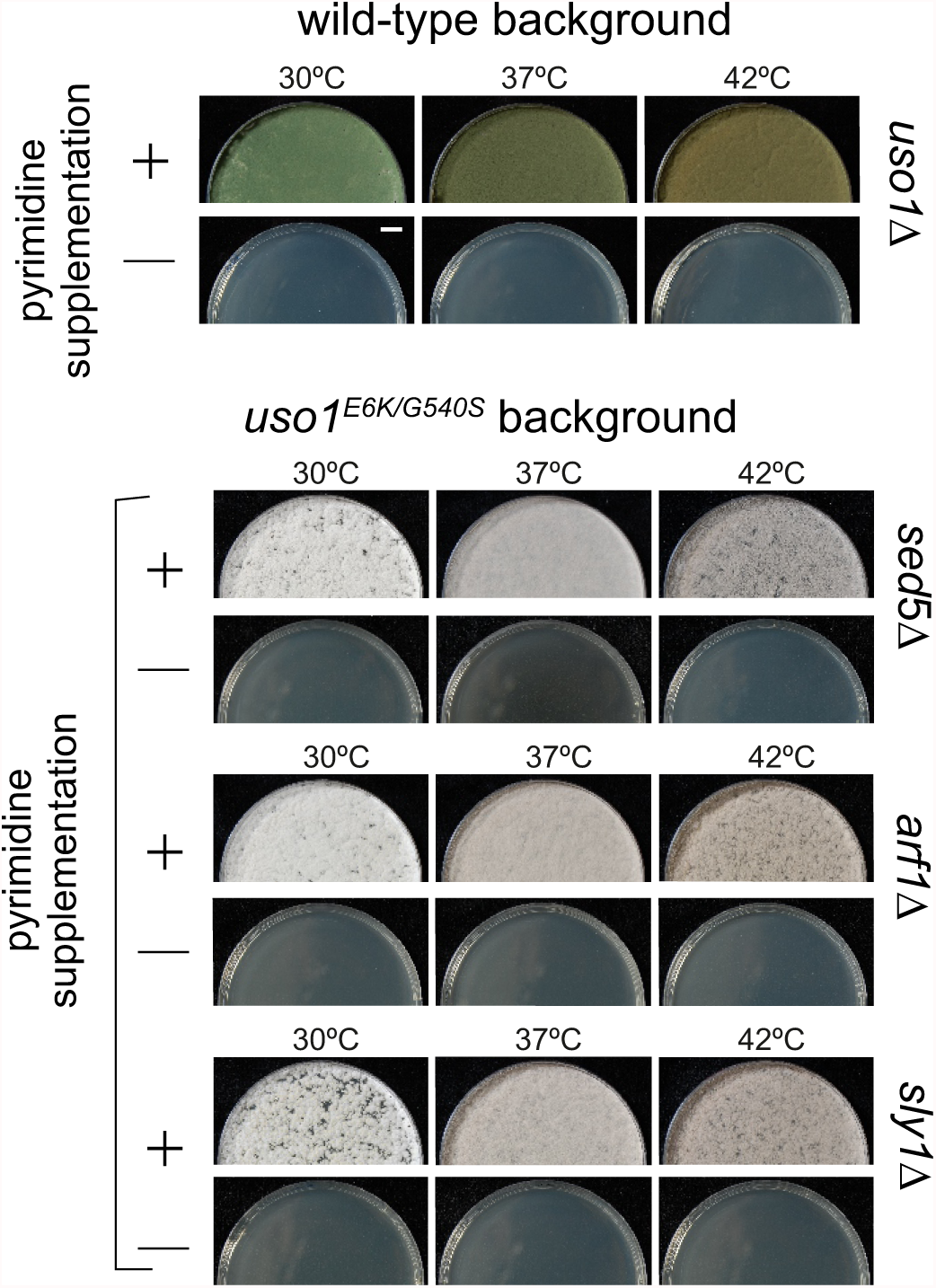
E6K/G540S do not rescue lethality resulting from *arf1*Δ, *sly1*Δ or *sed5*Δ. Top, *uso1* is an essential gene. Singly-nucleated conidiospores derived from a heterokaryotic strain in which one class of nuclei carries a deficient *pyrG* uracil biosynthetic gene whereas the second class contains a *uso1*Δ allele tagged with functional *pyrG* were unable to grow on medium lacking pyrimidines at any of the tested temperatures. Bottom: Similar experiments showing that unlike *rab1*Δ strains, strains carrying lethal *arf1*Δ, *sly1*Δ and *sed5*Δ alleles cannot be rescued by *uso1^E6K/G540S^*. Top panel, strains with green conidiospores; bottom, strains with white conidiospores.

**Figure 2—figure supplement 1.**
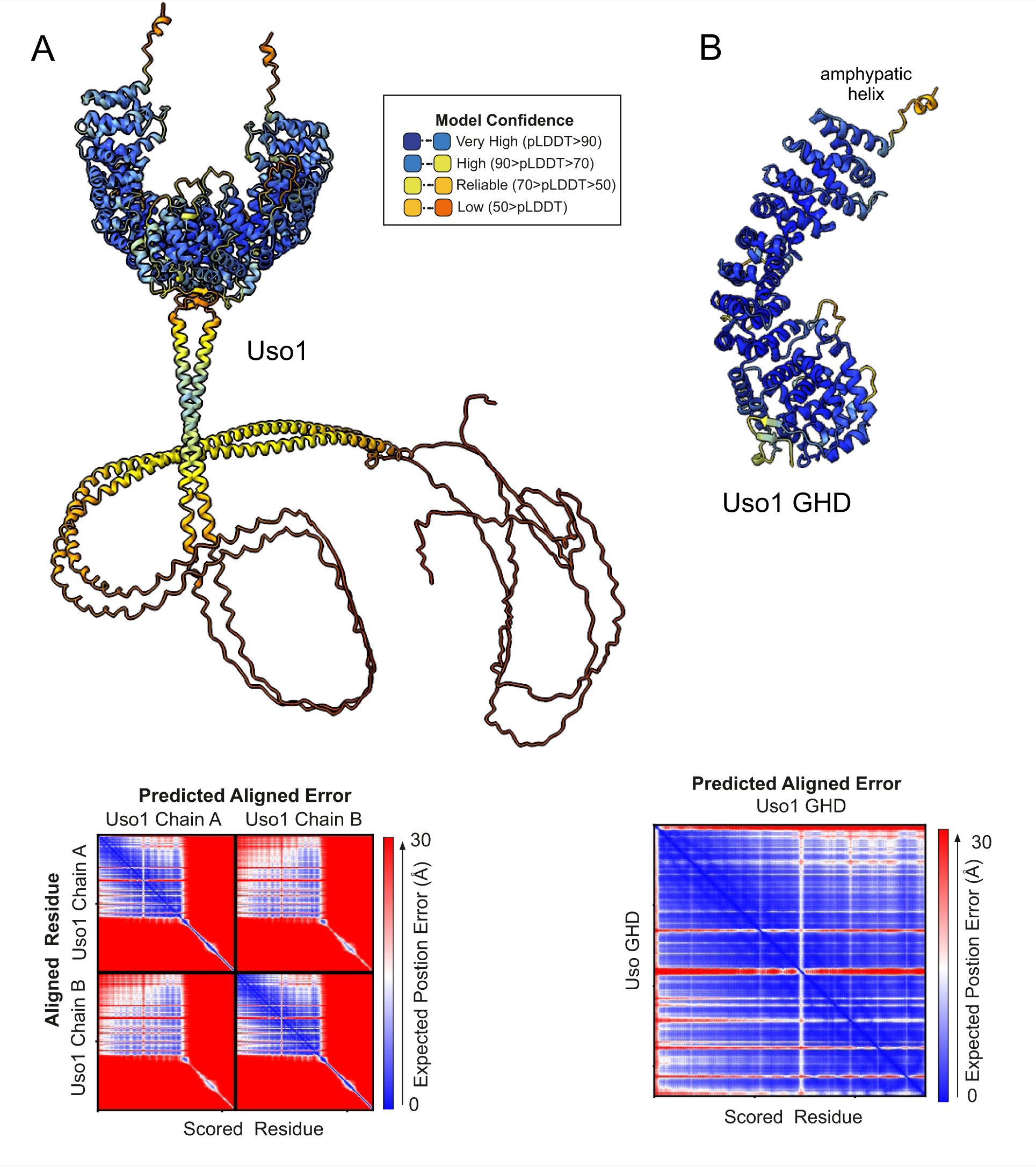
AlphaFold2 predictions of Uso1. Ribbon representation of AlphaFold 2-predicted structures of full-length Uso1 (A) and Uso1 GHD (B), color-coded by pLDDT values. Graphs at the bottom are the corresponding plots of predicted aligned error of the residues (PAE).

**Figure 3—figure supplement 1:**
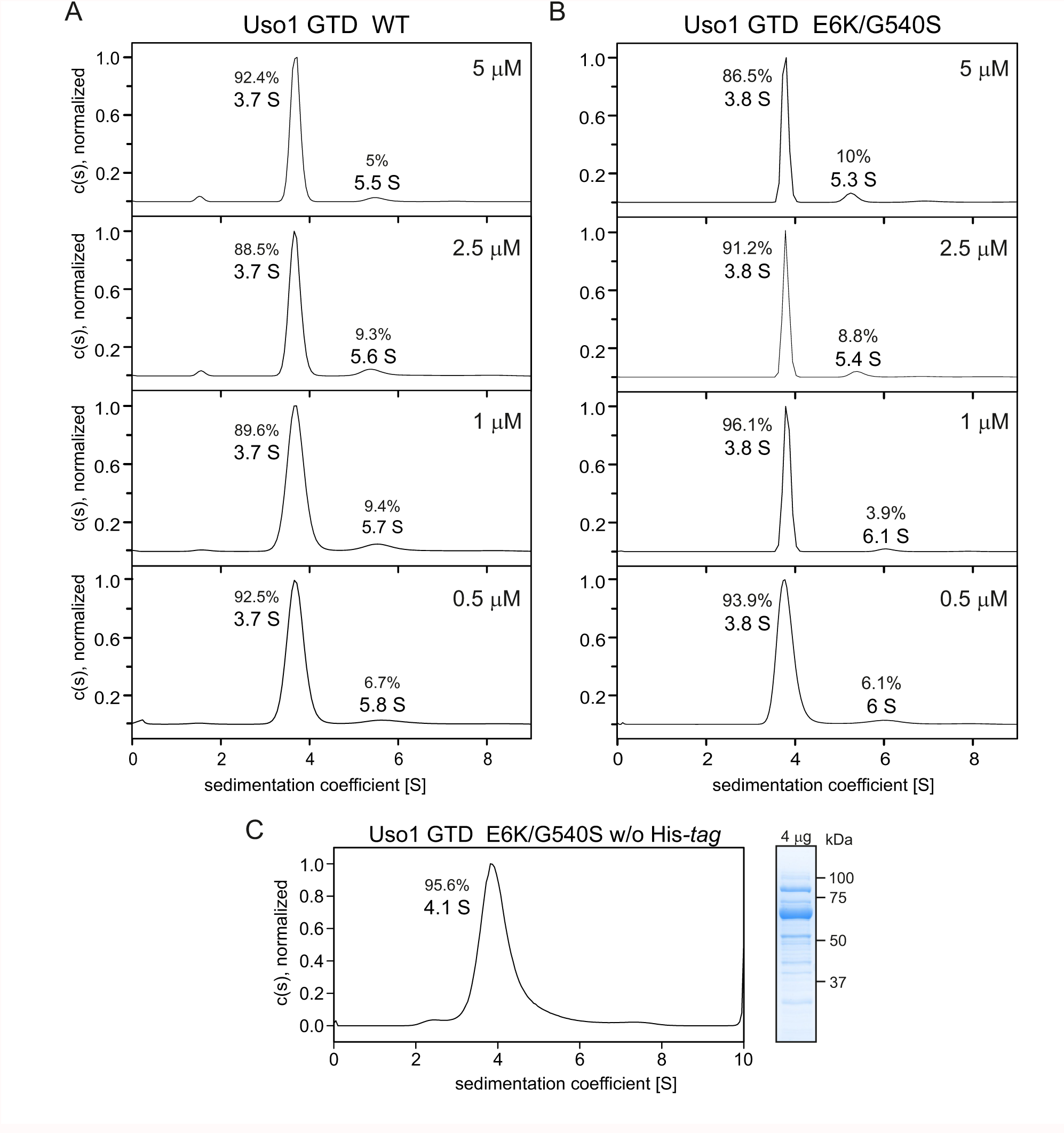
GHD is a monomer across a range of concentrations. (A) and (B). Sedimentation velocity experiments with wild-type and E6K/G540S mutant GHD, respectively, showing that they behave as monomers at concentrations up to 5 μM.(C). Sedimentation velocity profile of E6K/G540S mutant GHD lacking the His-tag, showing that the presence of the latter does not interfere with oligomerization, and a picture of a Coomassie stained gel showing the purity of the protein preparation on the right.

**Figure 4—figure supplement 1:**
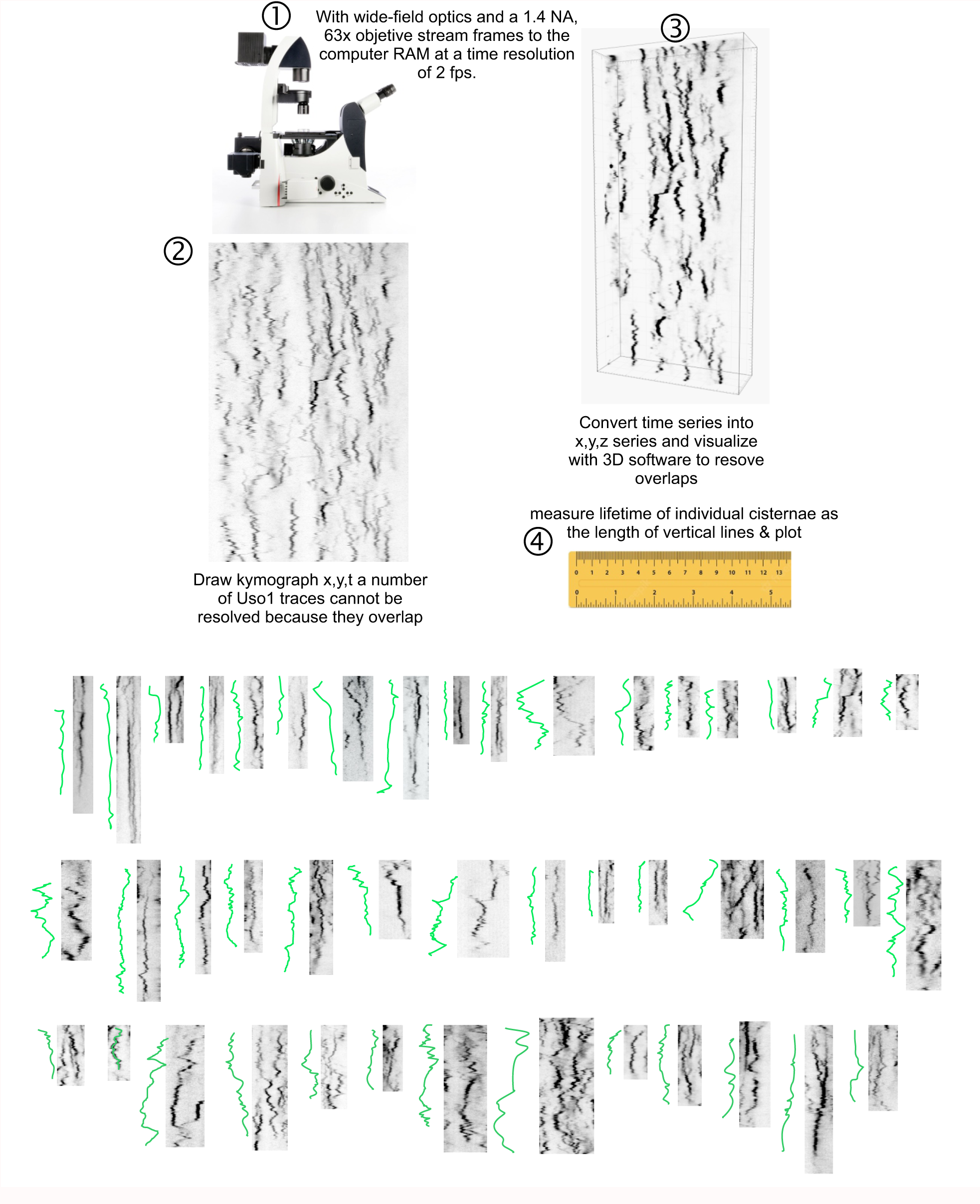
Methodology for tracking the half-life of Uso1-GFP on punctate structures. (1) 3D movies were acquired by streaming pictures to the computer RAM at 2 fps. Appropriate reduction of excitation light intensity permitted acquisition of 400 frames without apparent phototoxicity. (2) The behavior of punctate structures over time was represented in kymographs, in which vertical lines represent the time of residence of Uso1 on membranes. (3). As vertical lines frequently overlapped, jeopardizing the quality of this analysis, we imported the time series into a 3D viewer as if they were (x, y, z) series. Rotation across the different axes facilitated unambiguous tracking of the trajectories across time. (4) The length of the trajectories was measured and converted to time units. Bottom graphs display examples of time trajectories.

**Figure 8—figure supplement 1:**
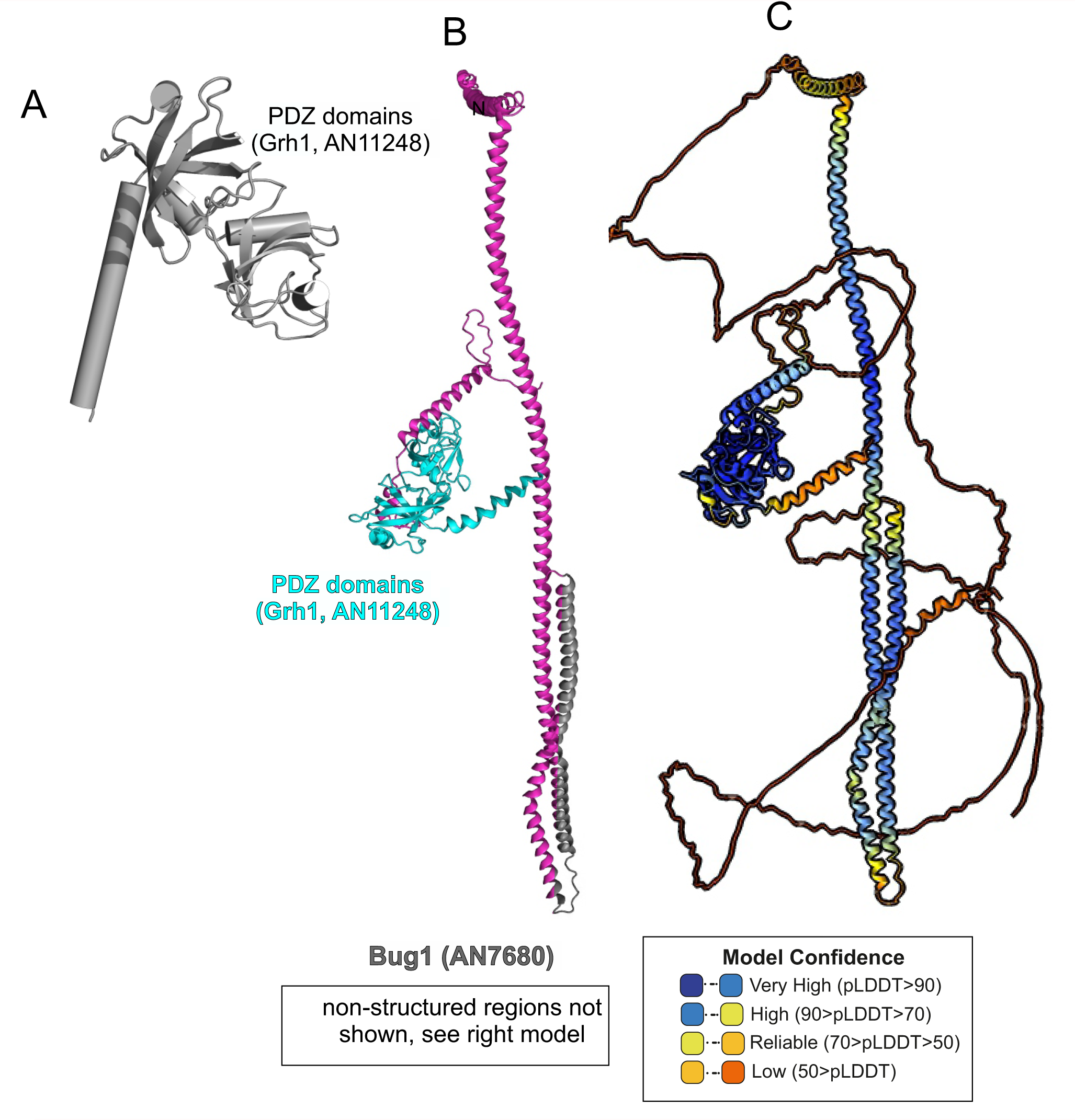
AlphaFold2 modelling of Grh1-Bug1. (A). Cartoon, with alpha-helices shown as cylinders, of the nearly N-terminal PDZ domains of Grh1 (B). AlphaFold 2 prediction of a 1:1 Grh1-Bug1 complex, trimmed of disordered regions (C). complete AlphaFold2 model of Grh1-Bug1 with color-coded model confidences values.

**Figure 10—figure supplement 1.**
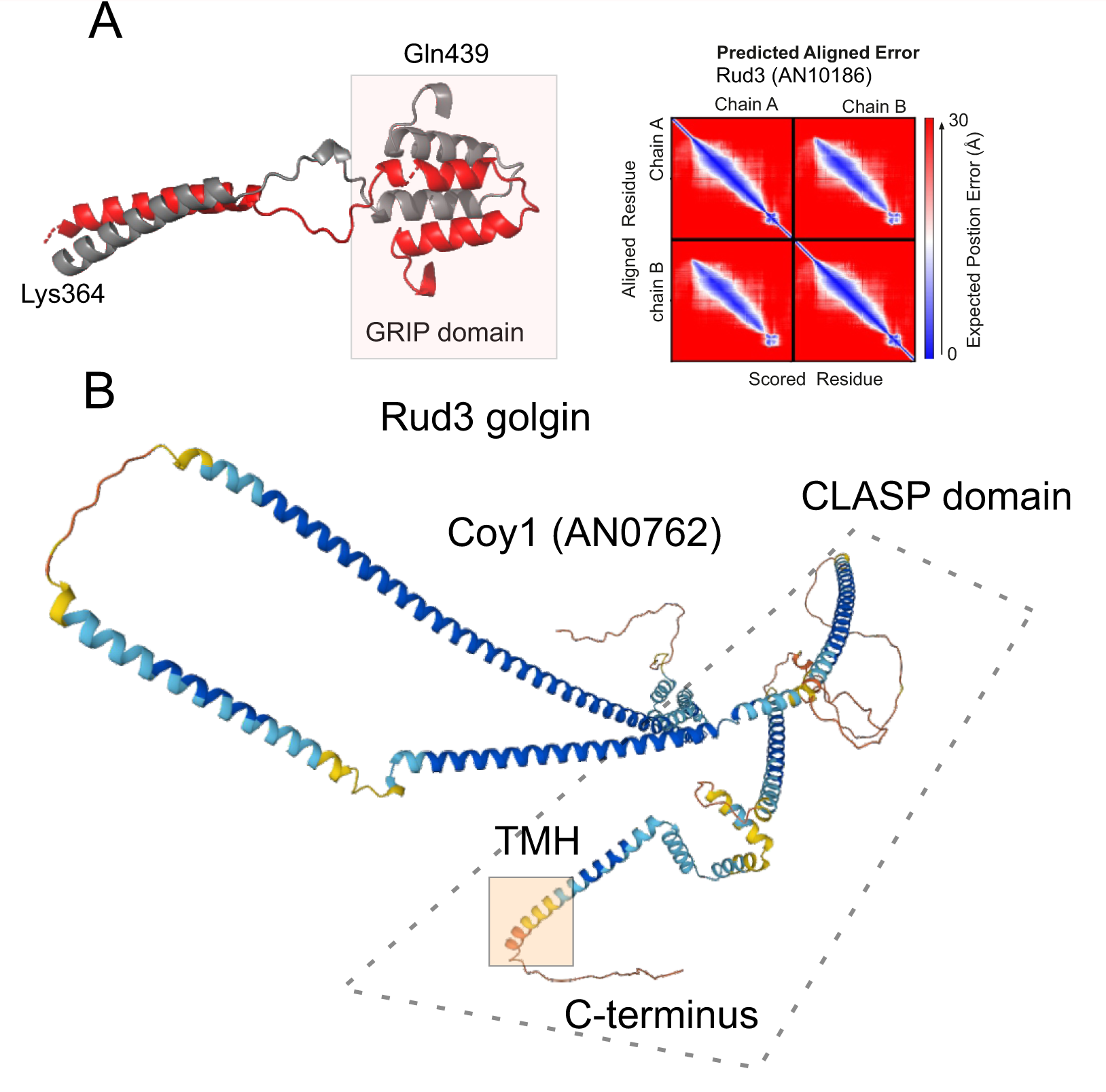
(A) AlphaFold2 model, with PAE plot, of the GRIP domain of A. nidulans RUD3, predicted to be a dimer. (B) AlphaFold2 model of Coy1, calculated as a monomer. TMH is the nearly C-terminal transmembrane helix that contributes to its recruitment to membranes

**Figure 10—figure supplement 2:**
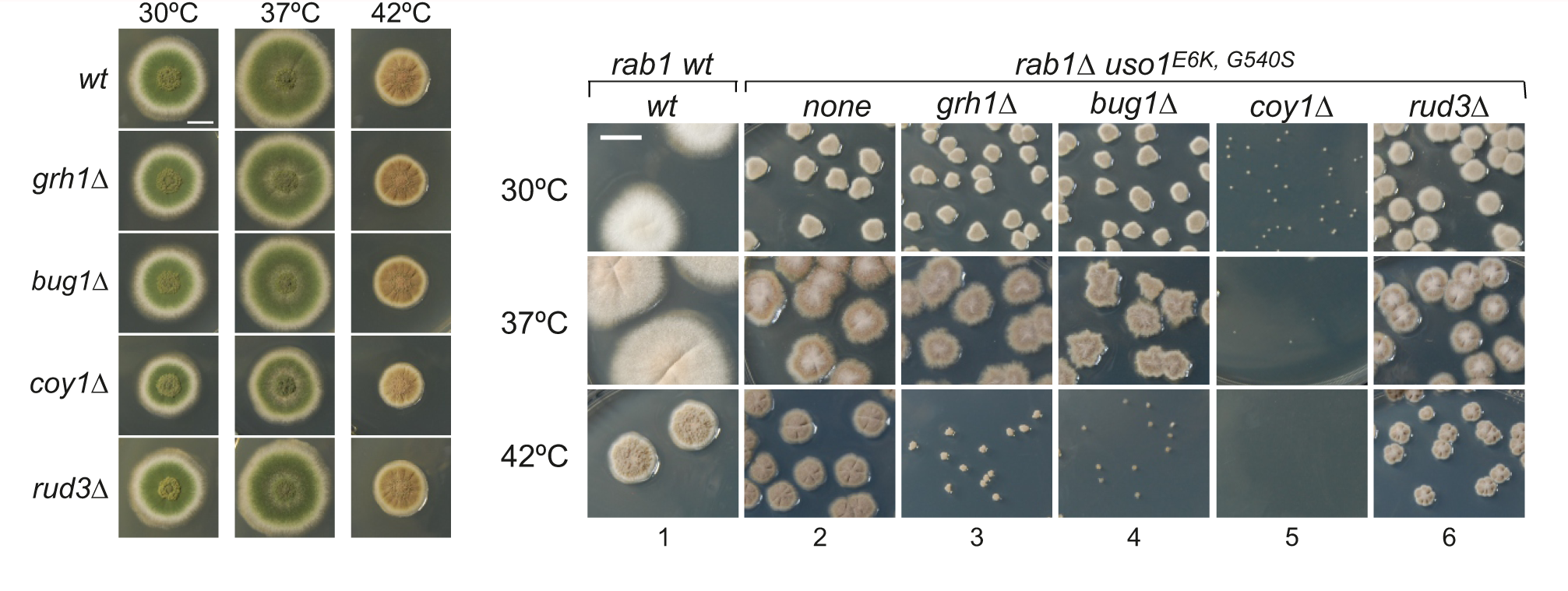
Growth phenotypes of null mutants of genes encoding golgins. (A). Ablation of individual golgins Bug1/Grh1 and Rud3 does not result in detectable growth defects. *coy1*Δ strains have a subtle growth phenotype. (B). Negative effects of *grh1*Δ, *bug1*Δ, coy1Δ and *rud3*Δ on the ability of *uso1^E6K/G540S^* to rescue *rab1*Δ. Note that *coy1*Δ and *rab1*Δ *uso1^E6K/G540S^* are synthetically lethal. For convenience, this set of strains carried a mutation resulting in white conidiospores, as opposed to the wild-type green color.

**Figure 11—figure supplement 1.**
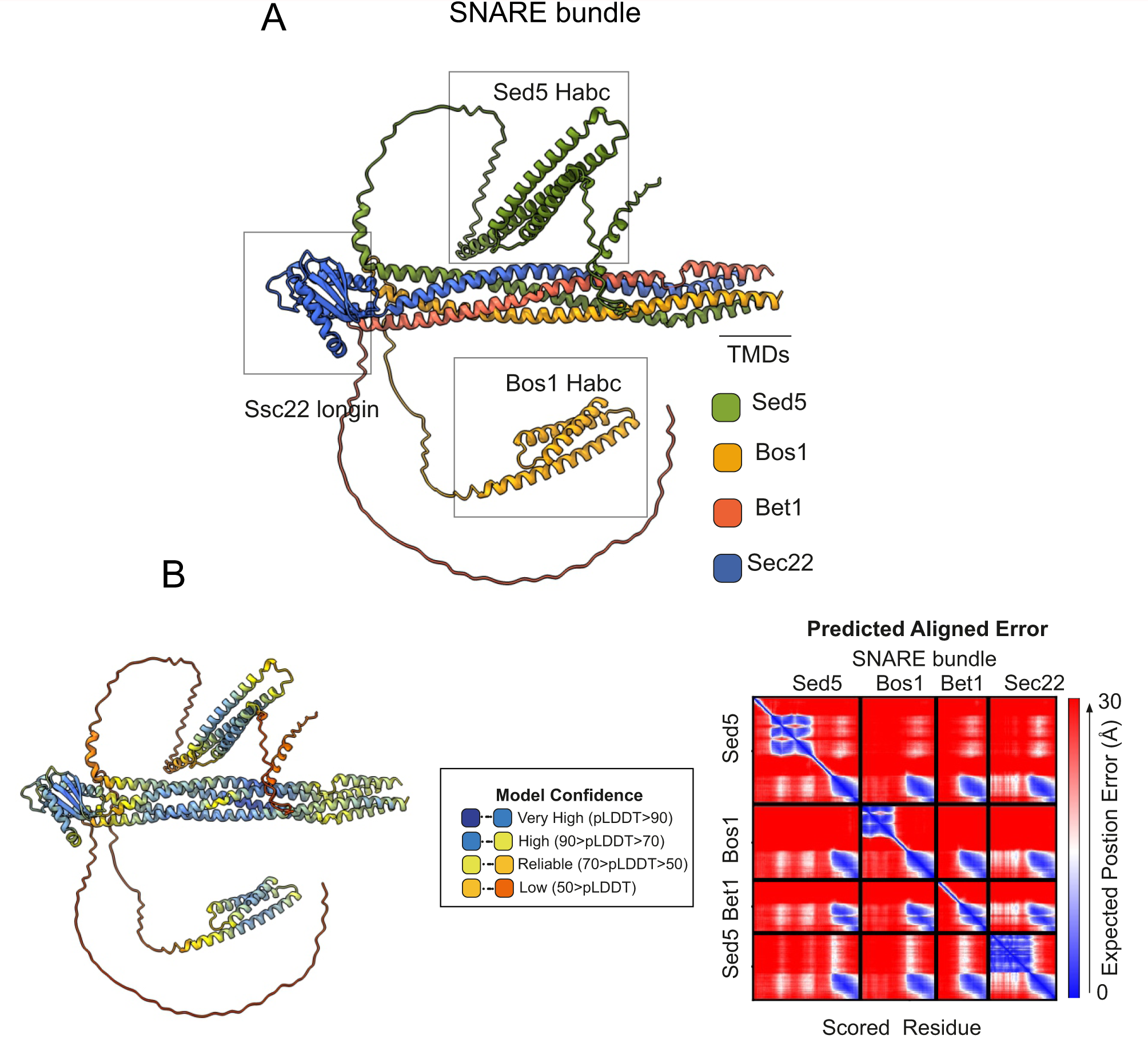
AlphaFold2 prediction of the ER/Golgi SNARE bundle. (A) Sec5/Bos1/Bet1/Sec22 predicted SNARE bundle. (B) Quality control (pLDDT, color coded, and PAE) of the model.

**Figure 12—figure supplement 1.**
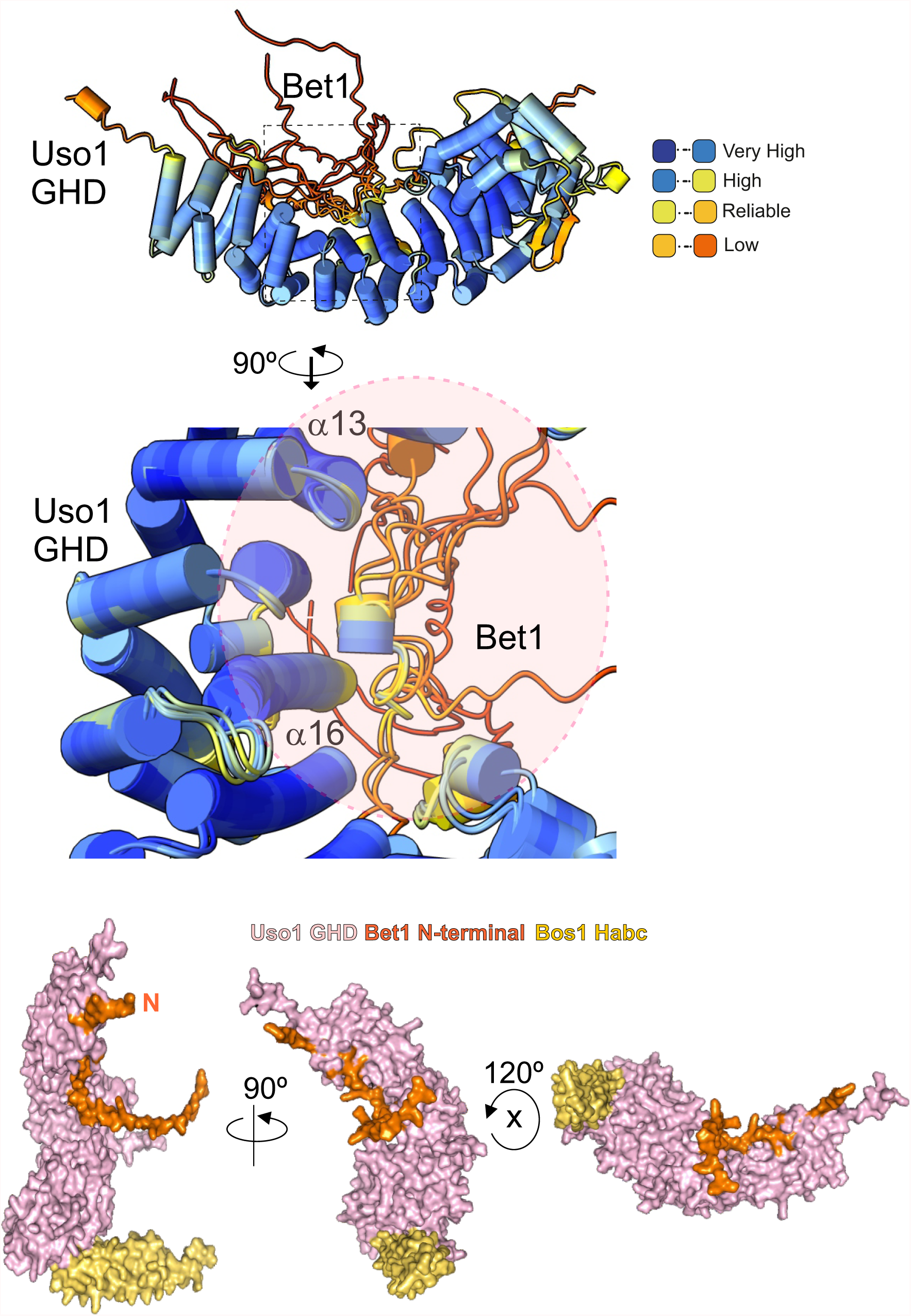
AlphaFold2 prediction of the Bet1-GHD interaction. The putative binding surface of Bet1 and Uso1 as determined by AlphaFold2. Top images, cartoon of Bet1-GHD interactions, colored by pLDDT score. Alignment of four independent predictions involving the Bet1 N-terminal region and Uso1 GHD. A single model for Uso1 GHD is shown on the top representation for simplicity. In spite of the disordered nature of the N-terminal Bet1 region, the Bet1-Uso1 binding interface is consistent among models. Bottom, surface representation of the N-terminal Bet1 region (orange) in complex with the GHD. Also indicated is the Habc domain of Bos1 (yellow) bound to the GHD.

**Figure 12—figure supplement 2.**
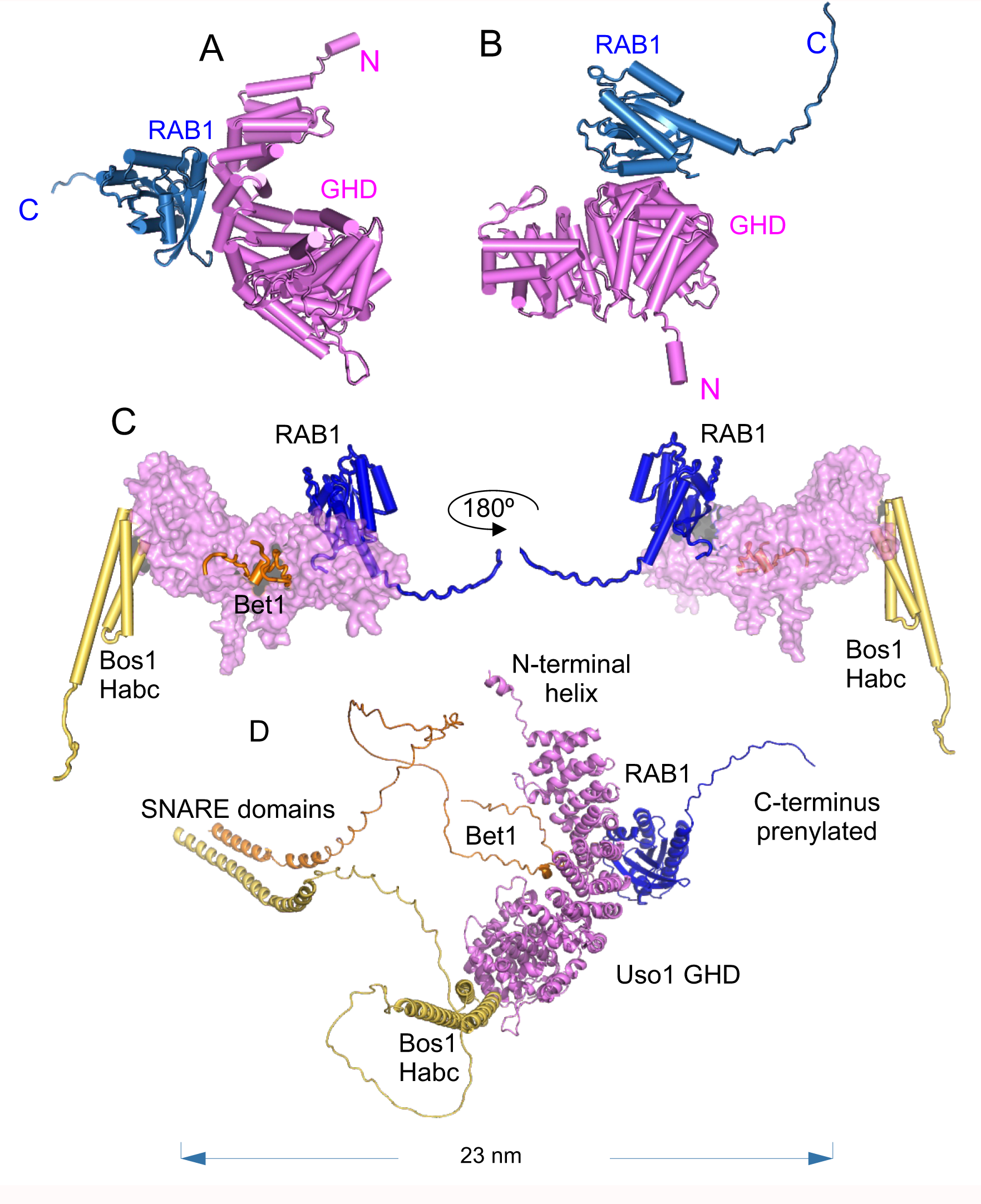
AlphaFold 2 prediction of the RAB1 binding site on the Bet1/Bos1/Uso1 GHD complex. (A) and (B): cartoon representation of the GHD-RAB1 complex. The model is depicted as pipes and planks (C): Orthogonal views of the Uso1 GHD-RAB1-Bet1-Bos1Habc structural model. The Uso1 GHD is shown as surface to emphasize the distant binding sites of the Bos1 Habc domain, RAB1 and the Bet1 N-terminal region (D): Ribbon representation of the model shown in C but including the full-length SNARE subunits, i.e. the GHD domain of Uso1, RAB1 and the SNARES Bet1 and Bos1. Proteins are in the correct orientation to connect membranes separated by 23 nm, counting from the SNARE TMDs to the prenylated RAB1 residues.

**Figure 12—figure supplement 3.**
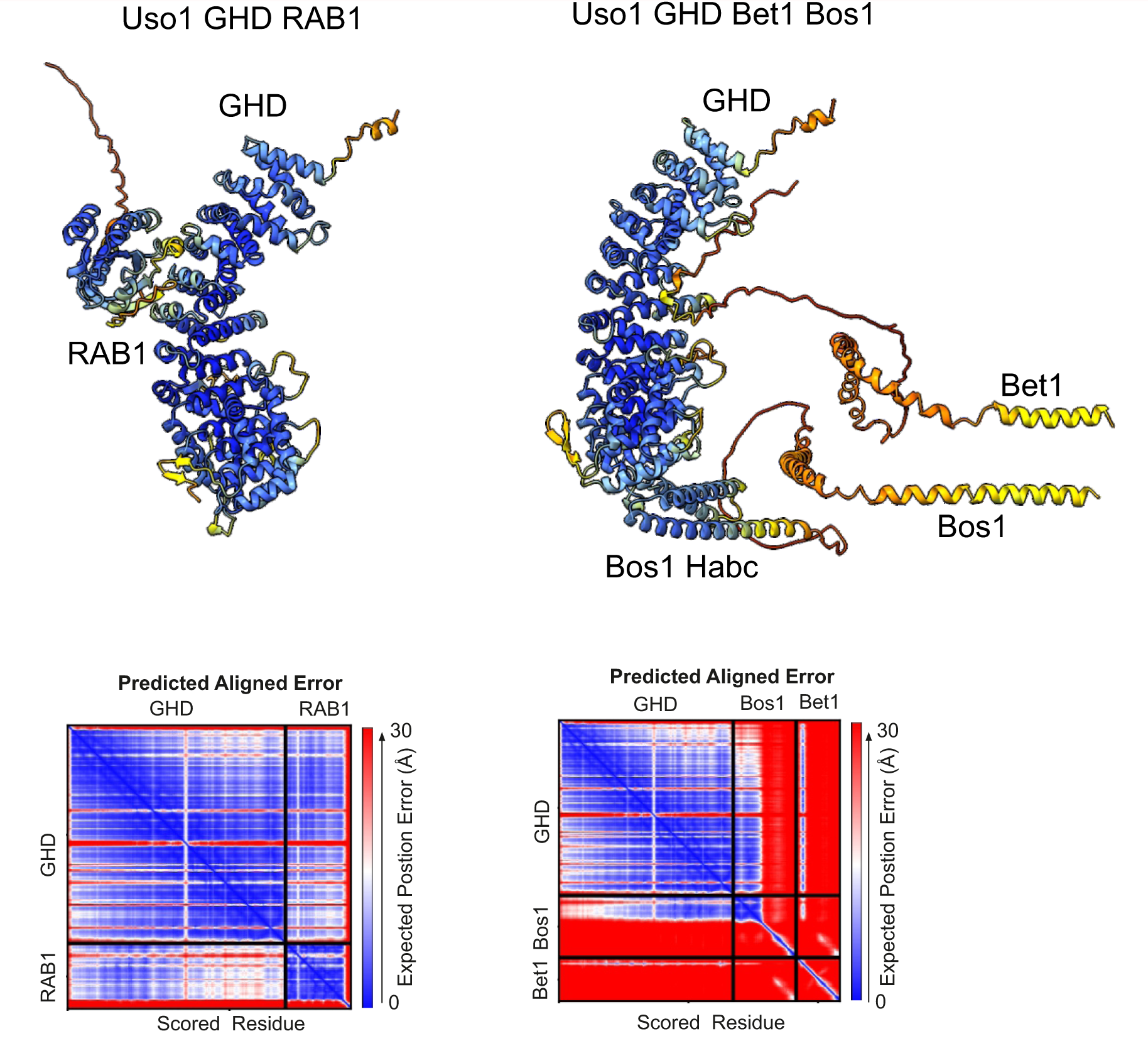
Quality control assessment of AlphaFold2 predictions for the indicated complexes.

**Supplemental Table I**

List and complete genotypes of *A. nidulans* strains used in this work

**Supplemental Table II**

Primers used for PCR-based genetic manipulations

**Figure 8 source data**

Raw images for western blots in panel D and uncropped pictures with used exposures and regions indicated.

**Figure 9 source data**

For panels B, C, D; raw images for western blots and uncropped pictures with used exposures and regions indicated.

**Figure 10 source data**

Raw images for western blots and silver-stained gels and uncropped pictures with used exposures and regions indicated.

**Figure 11 source data**

Raw images for western blots and Coomassie-stained gels and uncropped pictures with used exposures and regions indicated.

## Rich file media

**Video 1: Shaded 3D reconstruction of a hypha expressing Uso1-GFP Video 2: 4D acquisition showing the dynamics of Uso1-GFP.**

4D (x, y, z, t) in which Z-stacks were acquired at a rate of 1 frame every 2.6 sec

**Video 3: Dynamics of Uso1-GFP at 2 fps**

3D acquisition (200 frames) showing the dynamics of Uso1-GFP. Time resolution, 2 fps

**Video 4: Single Uso1-GFP cisterna tracked over time**

Example of Uso1-GFP cisterna. The video contains 96 photograms acquired at 2fps

**Video 5: ·3D reconstruction of a hypha expressing fluorescently labeled Uso1- GFP and Sec13-mCh**

There is little colocalization between Uso1-GFP and Sec13 ERES

**Video 6: ·4D video (1 fpm) of a hypha expressing fluorescently labeled Uso1-GFP and Sec7-mCh**

Uso1 does not colocalize at all with the TGN marker Sec7

